# A high-throughput 3D culture microfluidic platform for multi-parameter phenotypic and omics profiling of patient-derived organoids

**DOI:** 10.1101/2024.12.28.630600

**Authors:** Oronza A. Botrugno, Elena Bianchi, Jolanda M. Bruno, Giovanni F.M. Gallo, Claudia Felici, Eduardo Sommella, Paola De Stefano, Valentina Giansanti, Valeria Rossella, Fabrizio Merciai, Vicky Caponigro, Valentina Golino, Danila La Gioia, Antonio Malinconico, Emanuele Colombo, Giulio Giovannoni, Gabriele Metalli, Guido Gatti, Gazziero Virginia Anna, Silvia Buonanno, Eleonora Guidi, Giulio Caravagna, Federica Pedica, Francesca Ratti, Carlo Tacchetti, Dejan Lazarevic, Paola Falletta, Samuel Zambrano, Davide Mazza, Davide Cittaro, Roger D. Kamm, Pietro Campiglia, Giovanni Tonon, Gabriele Dubini

## Abstract

Patient-derived organoids (PDOs) are poised to become central tools both in clinical practice, to preemptively identify patient optimal treatments, and in drug discovery, overcoming the limitations of cancer cell lines. However, the use of PDOs in these settings has been hampered by several bottlenecks, including sample requirements, assay time, and handling in the context of high-throughput assays. We developed a <u>M</u>icrofluidic <u>P</u>latform for <u>O</u>rganoids culture (MPO) that miniaturises and simplifies PDOs cultures in a 384-plate format. Both retrospective and prospective clinical studies demonstrate MPO predictive value and the straightforward implementation in the clinical setting. MPO allows subcellular phenotypic screenings, as imaging-based applications like Cell Painting, target engagement analyses, alongside the comprehensive definition of PDOs’ genomic, transcriptomic, proteomic, lipidomic, and metabolomic landscapes. Harnessing the pleiotropic capabilities of MPO, we uncovered the role of EZH2 inhibitors to prevent the long-term emergence of resistance to RAS inhibitors in metastatic colon cancer. In all, we demonstrate the potential of MPO to impact clinical practice, alongside exploration of the mechanisms underlying compound response and resistance.

## Introduction

Knowing preemptively which therapy may benefit a specific cancer patient represents the most daunting unmet clinical need in oncology. This information could provide patients immediate relief from the disease while preventing unwanted toxicities, often severe in the case of cancer therapies. Insofar, the selection of the most appropriate treatment has been largely confined to targeted therapies, relying on the genomic analysis of patient-derived samples^1^. However, several large prospective clinical trials^2–5^, most prominently the NCI-MATCH trial^6,7^, have demonstrated a limited response rate in an intention-to-treat analysis for these genomic-based therapies, not reaching 5% of all cases^1^.

One enticing opportunity to overcome this sobering clinical reality is the combination of static genomic measurements with the direct testing of large arrays of compounds directly on patient-derived samples before administering any treatment, that is, the functional precision medicine (FPM) paradigm^1,8^. FPM efforts, proposed decades ago, have been hampered by several technological barriers, including rudimentary *ex vivo* culture methods and low-throughput capabilities, and thus they never entered clinical practice^9^.

Recent technological breakthroughs are starting to tackle these roadblocks^10^. Prominently, patient-derived organoids (PDOs), stem-cell derived self-organized 3D models, closely mimic *in vitro* the complex structure and function of the corresponding healthy or diseased *in vivo* tissue, including cellular heterogeneity, histopathological features, and in the case of cancer the genomic profiles of the corresponding tumour of origin^11^. They can be derived and swiftly expanded from the tissues of patients and stored in “living biobanks”^12^. In the last decade, PDOs have become a central tool for therapeutic screening and precision medicine, especially in the cancer field^13–16^, overcoming the limitations of standard two-dimensional (2D) cell cultures and even patients derived xenografts (PDX)^10,17^. Specifically, cells grown in monolayer, the primary model for drug screening, are easy and practical to culture but do not reproduce the complexity of a tumour tissue in terms of cellular heterogeneity, gene expression and multi-omic profiles, pathways activity, and drug sensitivity^18^. When compared with PDXs, PDOs are easier to work with, less time-consuming, more efficient, less expensive, and amenable to high-throughput drug screenings^10^. Notably, a tight correlation between patient and PDOs responses to drugs has been firmly established in several tumour types^14,16,19,20^. While medium-throughput drug screenings on PDOs have been proposed in retrospective models or to inform patients about second line treatments or adjuvant therapies, one of the major challenges and still unmet goals is to use PDOs as prospective models for defining the optimal treatments for cancer patients.

Crucial for the effectiveness of this approach is the development of innovative systems to perform high-throughput drug screens that reduce the time between PDOs derivation and testing, within a time frame that could inform clinical treatment decisions. Microfluidics may provide enticing opportunities for high-throughput screening (HTS). However, the need to use soluble Basement Membrane Extract (sBME) such as Matrigel™ and Cultrex™ to generate PDOs poses unique challenges, given sBME complex non-Newtonian rheology, irreversible thermal gelation, and low mechanical properties. Lowering the concentration of sBME down to 2 to 10% vol/vol^21^ certainly facilitates the use of robotics in HTS, but it remains problematic as it is suitable only for short-term cultures and does not fully recapitulate optimal organoid culturing conditions^21–23^. An additional challenge towards HTS implementation stems from the sedimentation of cell aggregates within the sBME, which results in heterogeneous seeding during the dispensing process.

Efforts are ongoing to complement traditional functional assays of cell viability, such as those based on intracellular enzyme activity or ATP, with methods that faithfully assess the cellular phenotypes emerging from exposure to drugs^24–28^. As a matter of fact, oftentimes effective therapeutic compounds do not elicit extensive cell death, nor select for genetic subclones, but exert more subtle and yet impactful dynamic phenotypic responses or mostly affect classes of molecules such as protein levels, post-translational modifications, lipids and metabolites, which are not captured by genomic or transcriptomic analyses^29,30^.

To implement the use of high-concentration sBME matrices in miniaturized formats for high-throughput drug screening, overcoming these rheological, construct sedimentation, and compartmentalization challenges, we engineered a <u>M</u>icrofluidic <u>P</u>latform for <u>O</u>rganoids culture (MPO), that miniaturises PDOs culture in extracellular matrix (ECM), enabling the determination of basic response parameters within a few days after the tissue explant. At the same time, MPO allows integrative multi-omics analysis and high-content imaging studies in PDOs-based HTS applications, including one of the most popular image-based assay for multiparametric cell profiling, Cell Painting^31^, as well as long-term PDOs cultures, thus addressing crucial questions such as the emergence of resistance to targeted therapies.

## Results

### MPO allows seeding and culturing of hundreds of PDOs

We have developed a microfluidic platform, MPO, through the combination of microfluidic capabilities with 3D printing approaches, that enables high-throughput seeding of matrix-embedded organoids, allowing culturing and drug testing of PDOs in a miniaturized format. MPO relies on a custom refrigerated microfluidic cartridge that maintains the bioink in a fluid phase, preventing premature gelation and ensuring consistent rheological properties during dispensing, regardless of the type or concentration of the matrix used (see Materials and Methods and ref.^32^) (Fig. 1a). To prevent axial sedimentation of organoids during dispensing, the cartridge tubing is arranged in a quasi-horizontal configuration and the flow is carefully regulated, resulting in homogenous seeding of matrix-embedded organoids among droplets. To support PDOs, we 3D printed microstructures, which we dubbed NESTs, serving as individual culture compartments. After the deposition of the PDOs into the NESTs, the whole structure, including the NESTs, is turned upside down and dipped over the 384-well plate (Fig. 1a and Supplementary Fig. 1a). NESTs support lateral diffusion of nutrients and drugs while maintaining physical separation between samples. This hang-drop configuration is compatible with standard 384-well plates, enabling the swift transfers of the entire PDOs set into new plates. Furthermore, this approach greatly simplifies media exchange and exposure to drugs and reagents without the disruptions associated with the handling of PDOs growing at the bottom of the plates. Importantly, this configuration enables miniaturized PDOs cultures to be preserved and tested for weeks, thereby avoiding the loss of material and preventing dehydration and contaminations between wells (Supplementary Fig. 1b and see below).

**Figure 1.**
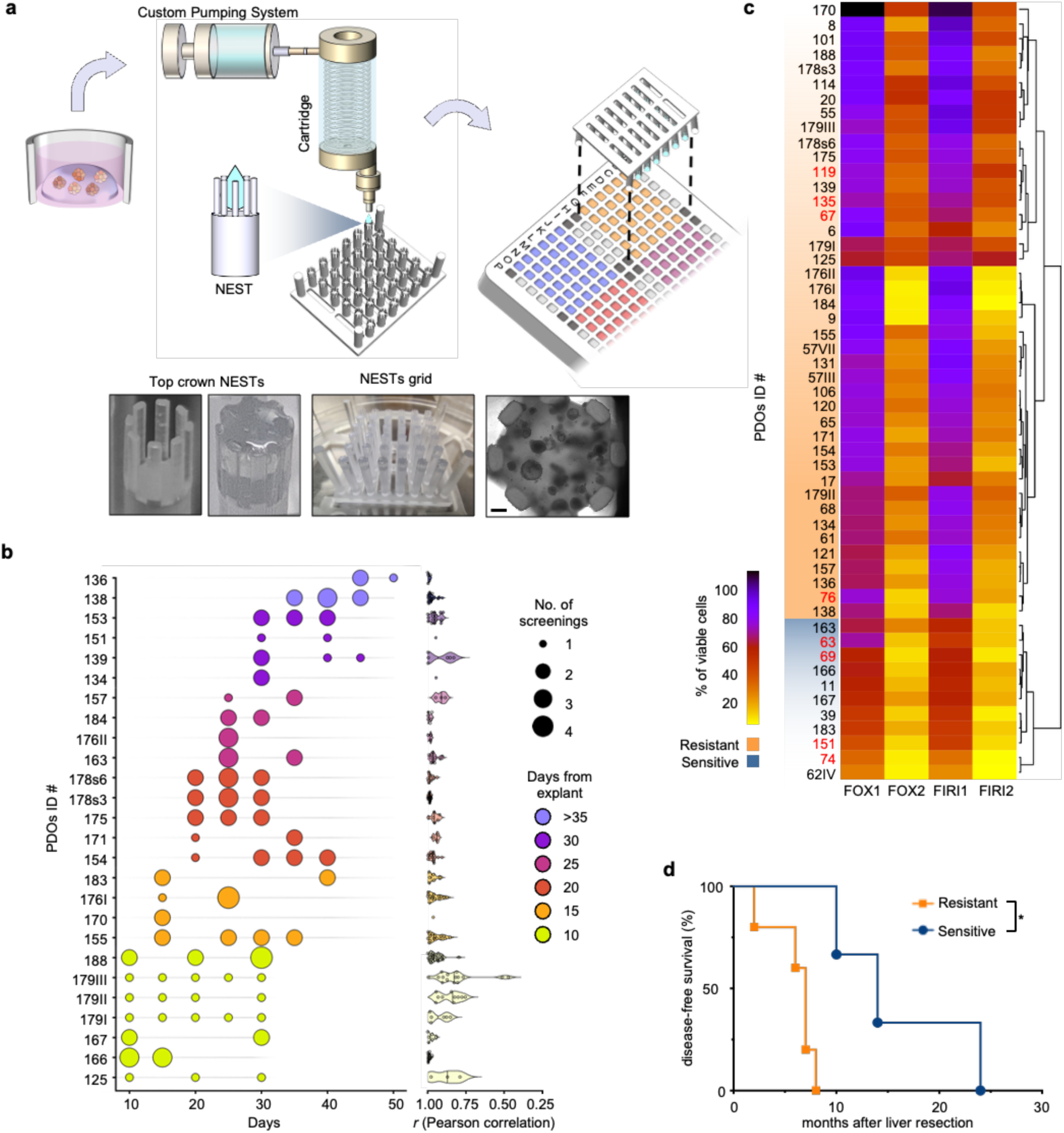
Drug sensitivity test on organoids grown and treated in the MPO. **a**. The MPO includes a pump, to dispense controlled volumes of samples, connected to a custom-made cartridge which enables the rapid and automated seeding of microsamples of organoids embedded in extracellular matrices. Microfluidic drops of the organoid suspension are seeded into individual NESTs organized in grids. The grids are then overturned and positioned in a 384-well plate, with each NEST dipped in the culture medium of a single well. The figure shows representative images of empty and filled top crown NESTs (on the left), a grid with the NESTs filled with the suspension of organoids before the culture (in the middle) and a brightfield image of organoids grown in the MPO (on the right). Scale bar: 200 μm. **b-c**. PDOs lines were seeded in the MPO and exposed to a combination of drugs that replicated two chemotherapeutic regimens commonly used to treat patients with mCRC, FOLFOX (FOX) and FOLFIRI (FIRI). Two concentrations were used, one highest (FOX2 and FIRI2) and one 10–fold lower (FOX1 and FIRI1). The sensitivities to the chemotherapeutic regimens was calculated as reported in Supplementary Figs. 2 and 3. At least two independent experiments were performed for each PDOs line. (b) The graph shows the results of a prospective study. It reports the time from explant to screening and the number of screenings performed at each time point, as well as the correlation between all screenings for each line (n= 26). (c) The heatmap represents the unsupervised clustering of the sensitivities to the chemotherapeutic regimens across all the PDOs lines included in the retrospective and prospective studies (n= 53). The range of responses among the different organoids allowed their separation into two major groups: sensitive and resistant. Based on the sensitivity to the lower doses, the threshold that distinguished the two groups resulted in a 40% reduction of viability relative to the untreated control (see Supplementary Figs. 2 and 3). In red organoids derived from patients receiving adjuvant chemotherapy after surgery. **d**. In the Kaplan-Meier curve, the organoids’ response data were matched with the clinical outcomes (reported as disease-free survival) of the corresponding patients if receiving adjuvant chemotherapy regimens based on 5-FU and oxaliplatin or 5-FU and irinotecan combinations (see Supplementary Fig. 1d). Gehan-Breslow-Wilcoxon (P=0.0244) and Log-rank (P=0.0134) tests were used to determine statistical significance among the two groups of patients (sensitive n= 3, resistant n= 5). *P < 0.05.

### MPO in the clinical setting

The pre-emptive identification of the optimal treatment remains the crucial unmet clinical need in oncology. The time required to obtain and test PDOs retrieved from patient samples remains a critical issue for the implementation of this technology in the clinical arena^10^. For example, in the SENSOR trial, the time for development of PDOs from metastatic colon cancer samples was 10 weeks^33^. We hence designed a prospective clinical study whose primary goal was the definition of the time needed after surgery to obtain and test PDOs in the MPO. We surmised that the small amount of material required by the MPO may shorten the time needed to obtain these data. The study, conducted from April 2023 to October 2024, included 60 patients undergoing surgical resections of metastatic colorectal cancer to the liver (mCRC), from which 66 tumour samples were collected and 37 PDOs derived, in line with previous publications, with the quality and size of resections, proportion of cancer cells and cell proliferation rate being the discriminating factors for the success of PDOs generation^10,14,16,33,34^. Of note, in almost 70% of the PDOs generated, we were able to complete multiple MPO drug testings within 40 days (Fig. 1b) (a clinically actionable time frame considering that the median time between surgery and the start of adjuvant therapy is 6 weeks^35^, with several cases screened a few days after sample retrieval (Fig. 1b).

We also sought to determine the MPO predictive value. To properly address this question, we first defined the threshold discriminating between sensitive versus resistant PDOs. To this end, we added to the prospective study a second retrospective cohort, for a total of 53 PDOs, exposed to a combination of drugs reproducing the two most common chemotherapeutic regimens used as first-line treatments for patients with mCRC, namely FOLFOX (FOX) and FOLFIRI (FIRI)^36^. Each chemotherapeutic combination was used at two doses, the maximum clinically achievable plasma concentration (Cmax) reported for each drug (FOX2 and FIRI2)^37–40^ and 10-fold lower (FOX1 and FIRI1), ensuring the exposure of PDOs at a range of drug concentrations relevant to those observed in clinical practice^41^. Drug effects were quantified after 96 hours of treatment using an ATP-based cell viability assay. Independent experiments demonstrated the robustness and reproducibility of the MPO (Fig. 1b, c and Supplementary Fig. 2, 3), also when compared with the conventional 96-well microplate setting (Supplementary Fig. 1c). Unsupervised clustering of the drug responses set the threshold distinguishing sensitive or resistant PDOs at 40% viability (Fig. 1c, Supplementary Fig. 2, 3). We hence relied on this threshold to determine the predictive value of MPO in a subset of patients for which the outcome after adjuvant chemotherapy was known (Supplementary Fig. 1d). Patients who were defined as resistant based on their PDOs response had a significantly shorter disease-free survival (DFS, 7 months), compared with the DFS (14 months) of patients for whom PDOs were labelled as sensitive (Fig. 1d and Supplementary Fig. 1d).

These data suggest robust predictive capabilities of the MPO for FOLFOX and FOLFIRI chemotherapy and emphasise the strong potential of the MPO as a tool for anticipating clinical responses to treatments.

### Target engagement assessment of drug responses in MPO-PDOs

Determining whether a drug is engaging with its target is pivotal in both addressing the efficacy of a treatment in a specific patient, as well as in drug discovery to define the spectrum of activity of compounds^42–44^. We hence explored the feasibility of measuring target engagement also in the MPO miniaturized setting. To this end, we adapted to the MPO the HTRF (<u>H</u>omogeneous <u>T</u>ime <u>R</u>esolved <u>F</u>luorescence)-based protocol^45^, which combines fluorescence resonance energy transfer technology (FRET) with time-resolved measurement (TR) (see Material and Methods and Supplementary Fig. 5a). We selected PDOs presenting mutations in the KRAS gene and exposed them to the pan-KRAS inhibitor BI-2865^46^. To assess target engagement, we assayed the phosphorylation status of ERK, a direct target of RAS, alongside ERK total levels (Fig. 2a and Supplementary Fig. 5b). HTRF adapted to MPO demonstrated that BI-2865 strongly inhibits ERK phosphorylation in the treated samples without affecting ERK total levels (Fig. 2a and Supplementary Fig. 5b). A conventional western blot analysis of the same PDOs grown and treated in standard assay conditions confirmed MAPK pathways modulation upon BI-2865 treatment, as seen in the HTRF-based assay (Fig. 2b and Supplementary Fig. 5c, d).

**Figure 2.**
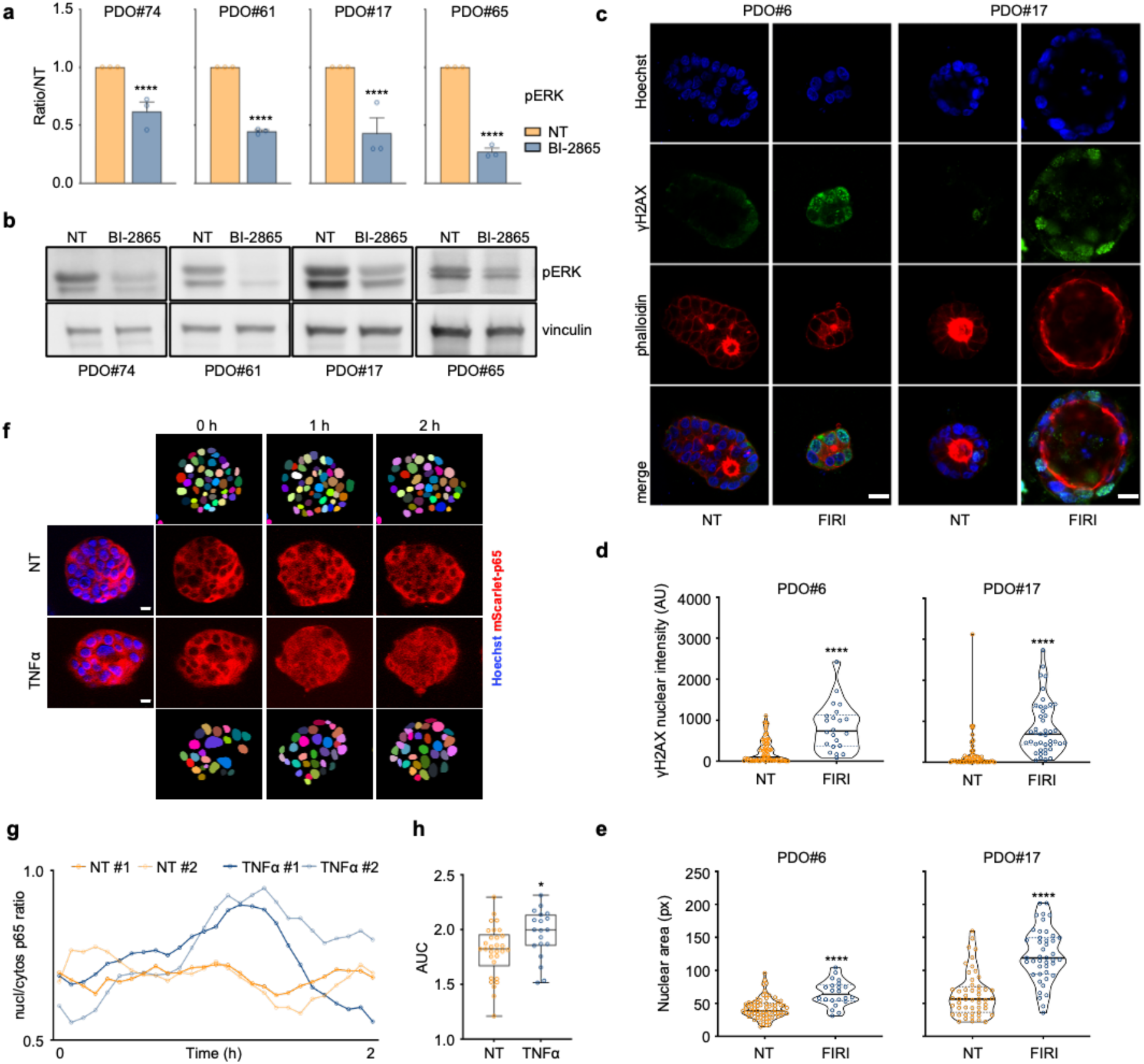
MPO allows for multi-parameter target engagement studies. **a**. A panel of four KRASmut PDOs lines were seeded in the MPO and treated for 4 hours either with DMSO (NT) or BI-2865 (KRASi). Levels of phospho-ERK (pERK) were assessed in HRTF-based assays. Results are reported as the levels of the protein in the treated samples relative to the untreated counterpart (ratio/NT) averaged (± s.e.m.) from three independent experiments, each performed in at least three technical replicates per condition. A two-way ANOVA followed by Sidak’s multiple comparisons test was used to determine statistical significance between groups. **** P<0.0001. **b**. Western blot analysis for the expression of pERK was performed on the same PDOs lines grown and treated as in (a) but not in a miniaturized setting. Vinculin was used as loading control. One representative experiment of two is shown. Quantification analysis for the assay is presented in Supplementary Fig. 5. **c-e**. PDO#6 and PDO#17 lines were seeded in the MPO and left untreated (NT) or treated with FOLFIRI (FIRI) for 96 hours. (c) Organoids were stained for Hoechst, Phalloidin, and γH2AX. Scale bar: 20 μm. Quantification of γH2AX nuclear intensity (AU, arbitrary units) (d) and nuclear area (px, pixels) (e) of NT (PDO#6 n= 72 and PDO#17 n= 51) and FOLFIRI-treated organoids (PDO#6 n= 22 and PDO#17 n=43). Statistical significance between groups was calculated by Bonferroni corrected Kolmorogov-Smirnov test. **** P<0.0001. **f-h**. MCF7 cells expressing mScarlet-p65 were grown as spheroids, seeded in the MPO and then stained with Hoechst. Spheroids were either left untreated (NT) or treated with TNFα and the p65 translocation into the nucleus followed by time-lapse for 2 hours. (f) Snapshots of the time-lapse images representative of NT and TNFα-treated spheroids under constant stimulation. For each group the masks used to segment and identify the nuclei are reported. Scale bar: 20 μm. (g) Representative single-cell profiles of nuclear to cytosolic m-Scarlet-p65 intensity for NT and TNFα-treated spheroids (two cells for each group). (h) Box plots show the quantification of the area under the curve (AUC) for the nuclear to cytosolic mScarlet-p65 intensity profiles quantified for cells in NT (n= 30) and TNFα-treated spheroids (n=19).

In all, these results demonstrate the feasibility and reliability of using MPO for high-throughput measurements of target engagement.

### High-content imaging of PDOs cultured in MPO

Imaging-based screening methods generate accurate measurements of multiple cellular features across conditions allowing phenotypic analyses, efficacy studies, target engagement and localization, thus providing crucial information on cellular states and transitions elicited by treatments. As such, this approach is relevant both for assessing the clinical response to drugs, and in HTS to obtain more reliable information on compound effects^26^. We hence tested whether imaging-based approaches could be applied also to the MPO. Mindful of the challenges posed by cells grown on a 3D system when compared with cells on 2D systems^47^, we used a modified version of the grid architecture, with a longer NESTs support. This modification positioned the surface of the matrix embedding PDOs closer to the bottom of the wells, reducing the working distance of the microscope objective and enabling the visualisation of the PDOs by confocal microscopy (Supplementary Fig. 6a).

We next assayed PDOs with quantitative immunofluorescence. To this end, using phosphorylated HA2X, a proxy of DNA damage^48^, we confirmed its nuclear staining after PDOs exposure to standard chemotherapies (Fig. 2c, d, Supplementary Fig. 6b and Supplementary movies 1, 2). Phalloidin staining of actin filaments confirmed that the treatment did not affect the overall morphological features of the PDOs (Fig. 2c). Of note, FOLFIRI treatment resulted in a significant increase in the nuclear area of both PDOs (Fig. 2e), a phenotypic modification already associated with chemotherapy exposure^49,50^.

We then determined if we could exploit MPO for 3D live-imaging. We recently engineered MCF7 cells, expressing a fluorescently tagged NF-kB mScarlet-p65^51^. We grew these cells as spheroids and transferred them to MPO. Similarly to what is observed in 2D cell cultures^51^, after stimulation with the NF-KB activator TNFα, we were able to detect the shuttling of mScarlet-p65 from the cytoplasm to the nucleus as early as 1 hour after stimulation (Fig. 2f, Supplementary Fig. 6c and Supplementary movies 3, 4). We next adapted our image analysis protocols^52^ to track single cells and quantify p65 activation by measuring the nuclear-to-cytosolic ratio of mScarlet-p65 intensity over time in both untreated and treated cells (Fig. 2g, h).

Overall these data demonstrate the suitability of the MPO to provide still and real-time imaging data.

### Cell Painting in the MPO-PDOs

We next explored whether we could harness other imaging methods not requiring the use of antibodies, nor relying on previous assumptions to explore and measure in a target-agnostic manner cellular responses to drugs or genetic manipulations. Cell Painting, one of the most prominent unbiased, quantitative, high-dimensional profiling techniques, relies on multiple fluorescent dyes to label organelles and cellular components, with the aim of capturing the phenotypic state of cells and their responses to perturbations^31^. This approach has been proven to be very effective in the context of 2D cellular models^31^, while several constraints have prevented its application in more complex model systems such as 3D organoids^53^.

We modified the 2D Cell Painting protocol to stain and image the 3D structures directly embedded in matrices. To test the staining, acquisition and analysis protocol, we used a six-dyes panel proposed by the Joint Undertaking Morphological Profiling (JUMP)-Cell Painting Consortium^54^ to comprehensively define eight subcellular components and organelles including nuclei, nucleoli and cytoplasmic RNA, f-actin, the Golgi apparatus and plasma membrane, mitochondria, and endoplasmic reticulum^47,55^ (Fig. 3a and Materials and Methods). We first exposed the reference 2D cell line U2OS^47^ to a selection of compounds^54^ proposed by the JUMP consortium that target relevant organelles and cell functions, along with the DMSO solvent control and confirmed the validity of the staining, and acquisition steps (Supplementary Fig. 7a). We then expanded our analysis to a PDOs line derived from a KRAS^G12A^-mutant tumor, which we treated also with FOLFOX, FOLFIRI as well as two KRAS inhibitors acting with via different mechanism of action: the pan-KRAS(OFF) inhibitor BI-2865^46^ and the RAS(ON) multi-selective inhibitor RMC-7977^56^. After acquisition of confocal microscopy images of U2OS cells (Supplementary Fig. 7a) and organoids at their maximal section focal plane (Fig. 3a, b), we designed for the analysis of both a custom pipeline in CellProfiler to segment individual cells into three compartments: nucleus, cytosol and whole cell (Fig. 3a, c and Supplementary Fig. 7a). We then extracted phenotypic features related to the single compartments that capture thousands of morphological characteristics (Supplementary Table 1). A machine learning algorithm then computationally defined a subset of 134 non-redundant features and identified similarities and phenotypic relationships elicited by the various drugs (Fig. 3d, Supplementary Fig. 7b, 8 and Supplementary Table1). Phenotypic profiles of PDOs treated with Cytochalasin B and Berberine localized at the edges of the biplot capturing all treatments in the phenotypic space (Fig. 3d), suggesting that in this model these treatments induced the most divergent phenotypes. Remarkably, the two 5-FU-based chemotherapies as well as the two KRAS inhibitors clustered together in the biplot, respectively. Clustering was similar for the same compounds at 24 and 48 hours, although more pronounced after longer treatments (Supplementary Fig. 8b). Additionally, when exploring the relationship between treatments and features, we observed that no individual feature single-handedly drove any of the morphological profiles induced by our panel of drugs, but rather cellular responses to each drug often spanned multiple channels^57,58^ (Fig. 3e). In particular, a very pronounced activity profile for the cytoskeleton and mitochondria channels was evident after the treatments with FOLFOX, FOLFIRI, and the RAS inhibitors (Fig. 3e, Supplementary Fig. 8a and Supplementary Table1), outcomes consistent with their known mechanisms of action and cytotoxicities^57,58^ (Fig. 3b, e).

**Figure 3.**
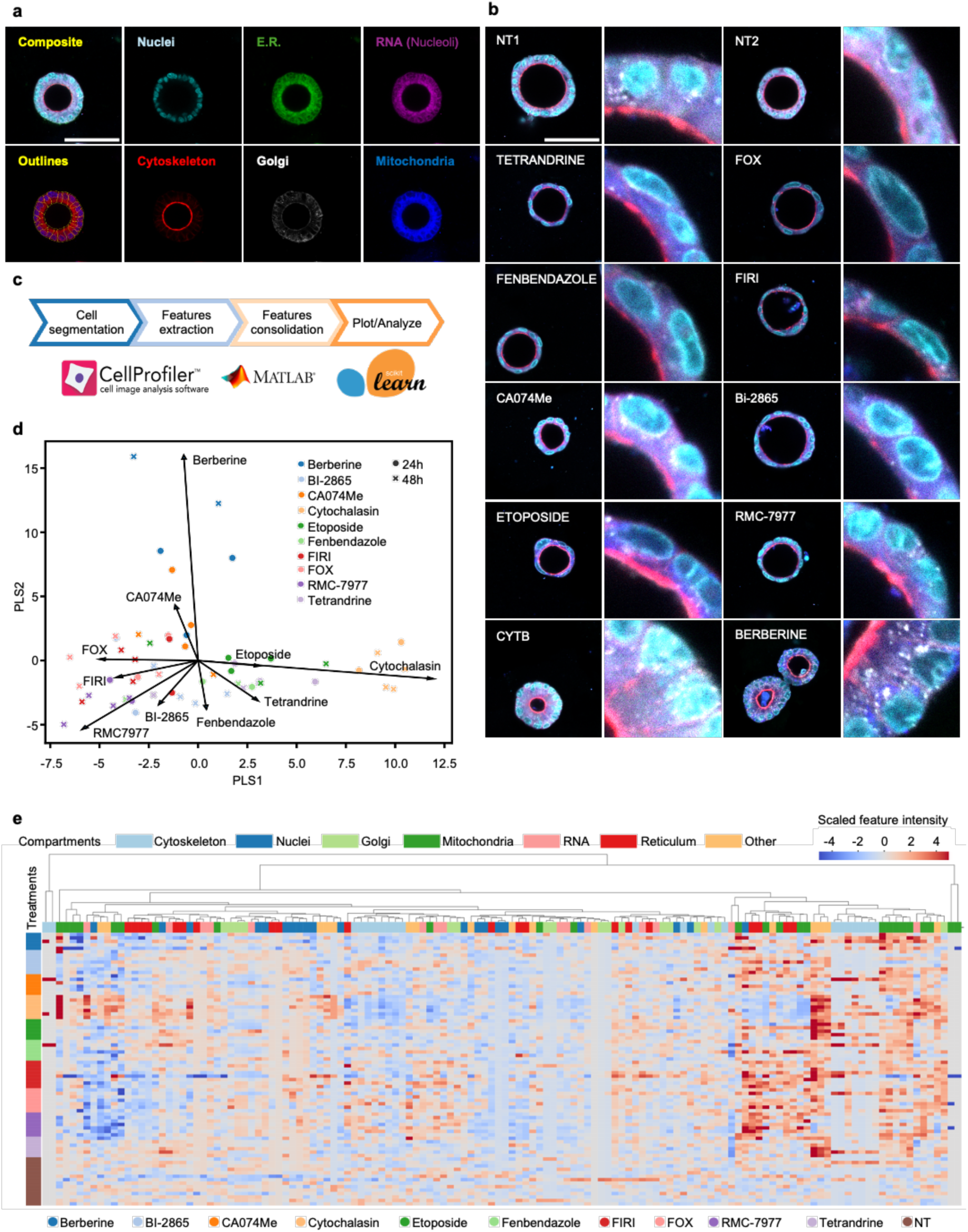
Cell Painting on organoids grown and treated in the MPO. PDO#17 was seeded in the MPO and left untreated (NT) or treated for 24 and 48 hours with Tetrandrine, Fenbendazole, Ca074Me, Etoposide, Cytochalasin B (CYTB), Berberine, FOLFOX (FOX), FOLFIRI (FIRI), BI-2865 and RMC-7977. **a**. Individual fluorescent channels from a representative single-plane confocal image acquired at the focal plane of the maximum organoid section. Composite image of all channels and segmentation-derived outlines are shown. Scale bar: 100 μm. **b**. Representative images of 48 hours-treated PDOs. NT1 refers to the vehicle-treated control in relation to the images of Tetrandrine, Fenbendazole, CA074Me and Etoposide samples and NT2 to the remaining treatments. Scale bar: 100 μm. **c**. Cell Painting images analysis pipeline. CellProfiler was used to segment cells and extract features, downstream analysis of features was performed with Matlab software and scikit-learn Python module. **d.** The biplot visualizes the profiles of multiple Cell Painting experiments in the PLS (Partial Least Squares) space fitted to maximize the differences between treatments. The arrows represent the loading vectors for each drug and indicate the relative contribution to each PLS coordinate displayed. **e.** The heatmap shows the scaled value of 134 non redundant Cell Painting features used in the analysis. Rows are sorted by treatment, columns are clustered according to the similarity of profiles across treatments. Column color code represents the different compartment grasped by each Cell Painting feature.

In all, these results demonstrate that MPO is amenable to multiparameter imaging-based phenotypic profiling, which complements traditional functional assays and provides a comprehensive depiction of cellular events that may reveal hidden vulnerabilities triggered by drug exposure.

### Genomic and transcriptomic profiling in the MPO-PDOs

The definition of the genetic lesions affecting a cancer and the identification of mutated or translocated proteins remain the mainstay of personalized medicine. Importantly, several reports have highlighted how mutated genes are not necessarily expressed in the corresponding cancer cells^59–62^, suggesting that it is imperative to determine not only the mutational status of a gene, but also whether it is expressed, to guide the selection of the most appropriate targeted therapies. To this end, we optimized the G&T-seq protocol^63^ to enable concurrent genome and transcriptome sequencing, thereby facilitating the integrated analysis of genetic variation and gene expression profiles from PDOs cultured on a single pillar. Whole genome sequencing (WGS) at low coverage on PDOs grown in the MPO showed that the copy number variants (CNV) profiles were very similar to those of PDOs grown conventionally and sequenced at high coverage (Fig. 4a). Then, we explored the feasibility of performing RNA-seq on the same PDOs within the MPO and to obtain insights on the response to treatments. To this end, PDOs were left untreated or exposed to the aforementioned chemotherapies, FOLFOX and FOLFIRI, and RNA was collected at 24 and 72 hours after treatment. RNA-seq analysis revealed clustering of PDO#61 and PDO#74 driven by identity. (Fig. 4b). Moreover, treated and non-treated PDOs were neatly separated based on the type of treatment and the separation was more pronounced after longer treatments (Fig. 4b).

**Figure 4.**
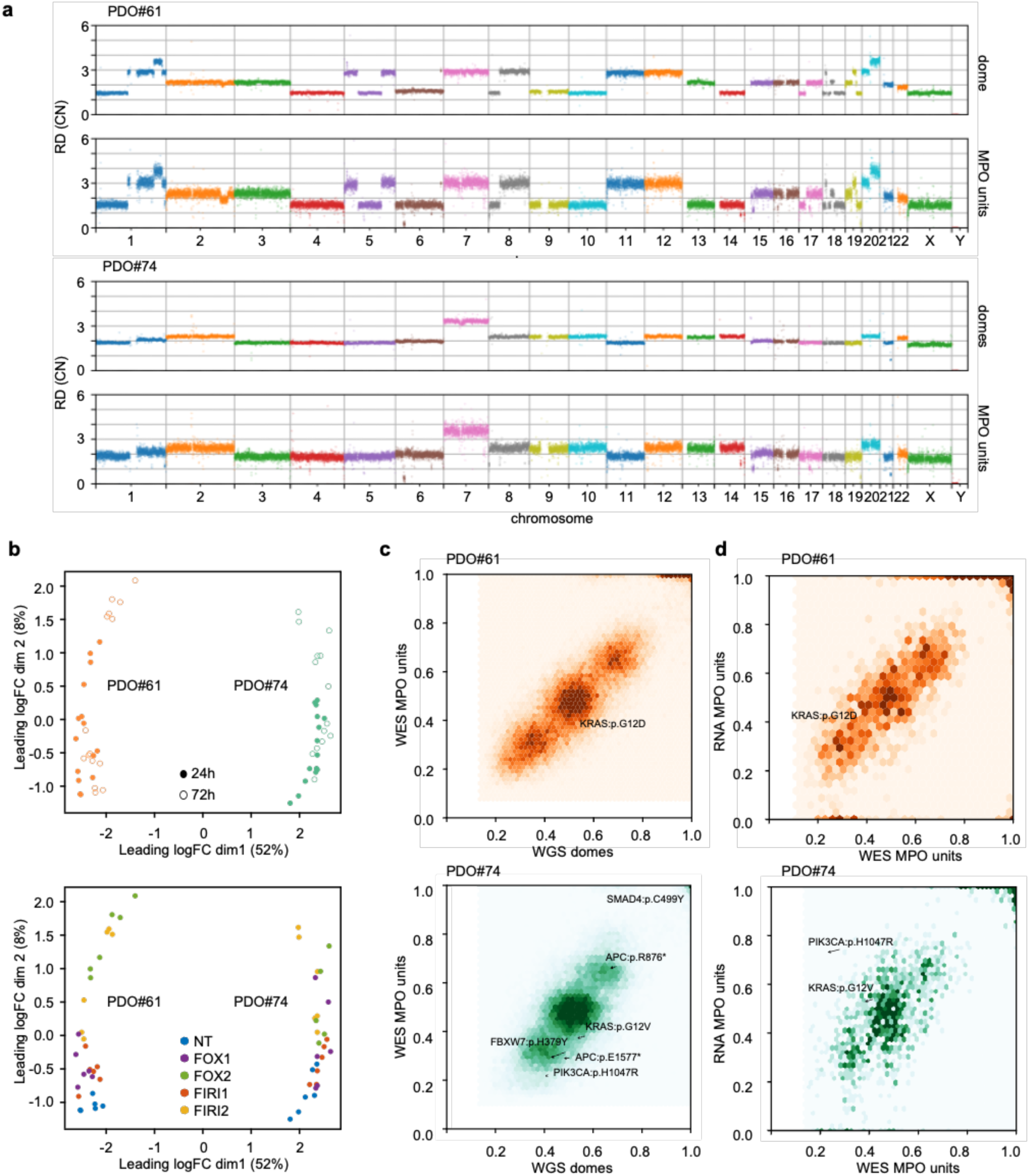
MPO enables the integration of multiple genomics platforms. PDO#61 and PDO#74 were seeded in the MPO and left untreated (NT) or exposed to FOLFOX (FOX) and FOLFIRI (FIRI), used at two concentrations, one highest (FOX2 and FIRI2) and one 10–fold lower (FOX1 and FIRI1), for 24 and 72 hours. **a.** Manhattan plot of the Read Depth (RD) signal for WGS performed on untreated PDO#61 and PDO#74 grown in the conventional setting (domes) and within the MPO. **b.** Multi-dimensional scaling (MDS) plots of RNA-seq expression profiles in two dimensions showing variation among samples. Axes indicate the percentage of variance explained by the corresponding MDS dimension. Each dot represents one sample. Samples are grouped by PDOs, with distinct clusters within each line representing the time of drug administration and treatment. **c.** Hexbin-plots showing comparative Variant Allele Frequencies (VAFs) between high coverage WGS and WES and **d.** WES *vs* RNA analysis for mutations in PDO#61 and PDO#74 on the MPO. Driver mutations are highlighted.

The reliability of the single-pillar G&T-seq approach was further supported by whole exome sequencing (WES) analysis on pillars. Compared to WGS, WES can provide better depth of coverage over targeted regions of interest coupled with easier data analysis, lower cost per sample and lower storage requirements^64,65^. Indeed, genetic variants detected in WES showed a high degree of concordance with variants detected in the WGS high coverage performed on PDOs grown in domes (Fig. 4c). To investigate if variants called from RNA-seq are sufficiently informative to depict the mutational status of the samples, we matched WES and RNA-seq data of untreated samples (Fig. 4d). While, as expected, not all cancer drivers genes were retrieved from RNA-seq data^59–62^, nonetheless there was a good concordance between WES and RNA data, with some intriguing expectations, such as the predominant expression of mutated PI3KCA in PDO#74 in line with previous reports^66^. The same results were obtained comparing RNA-seq to high-coverage WGS (Supplementary Fig. 9).

Overall, these findings support the possibility to perform reliable genomic and transcriptomic analyses within the MPO.

### Modelling drug resistance in PDOs cultured in the MPO

The emergence of resistance to anticancer therapies is one of the greatest challenges faced by clinical oncologists, especially when treating patients with targeted therapies. Given the multifaceted analyses and importantly, the long-term cultures that the MPO can support, we investigated whether we could use it to model the emergence of therapy resistance and characterize putative genetic and non-genetic driver mechanisms^67^. We exposed the *KRAS*^G12A^-mutant PDO#17 to the pan-KRAS(OFF) inhibitor BI-2865^46^ and the RAS(ON) multi-selective inhibitor RMC-7977^56^ at two different concentrations, or combined the two compounds. Concurrently using KRAS(OFF) and RAS(ON) inhibitors should interfere with two compensatory responses driving resistance to KRAS inhibition, the increased KRAS-GTP loading and the feedback activation of the RAS pathway^56,68–70^. The PDOs line was grown in the MPO under the constant presence of drugs for 25 days. While single treatments at lower concentrations were only initially effective, combination of both treatments showed synergistic activity compared to single agents alone, providing further evidence to support the hypothesis that combining RAS(OFF) and RAS(ON) inhibitors is beneficial in the context of adaptive resistance to KRASi^69^ (Fig. 5a). However, even the combination failed to quench cancer cell growth at later times. Intriguingly, treatment with RMC-7977 at the highest concentration prolonged growth inhibition, albeit at the end resistance emerged also during this treatment (Fig. 5a).

**Figure 5.**
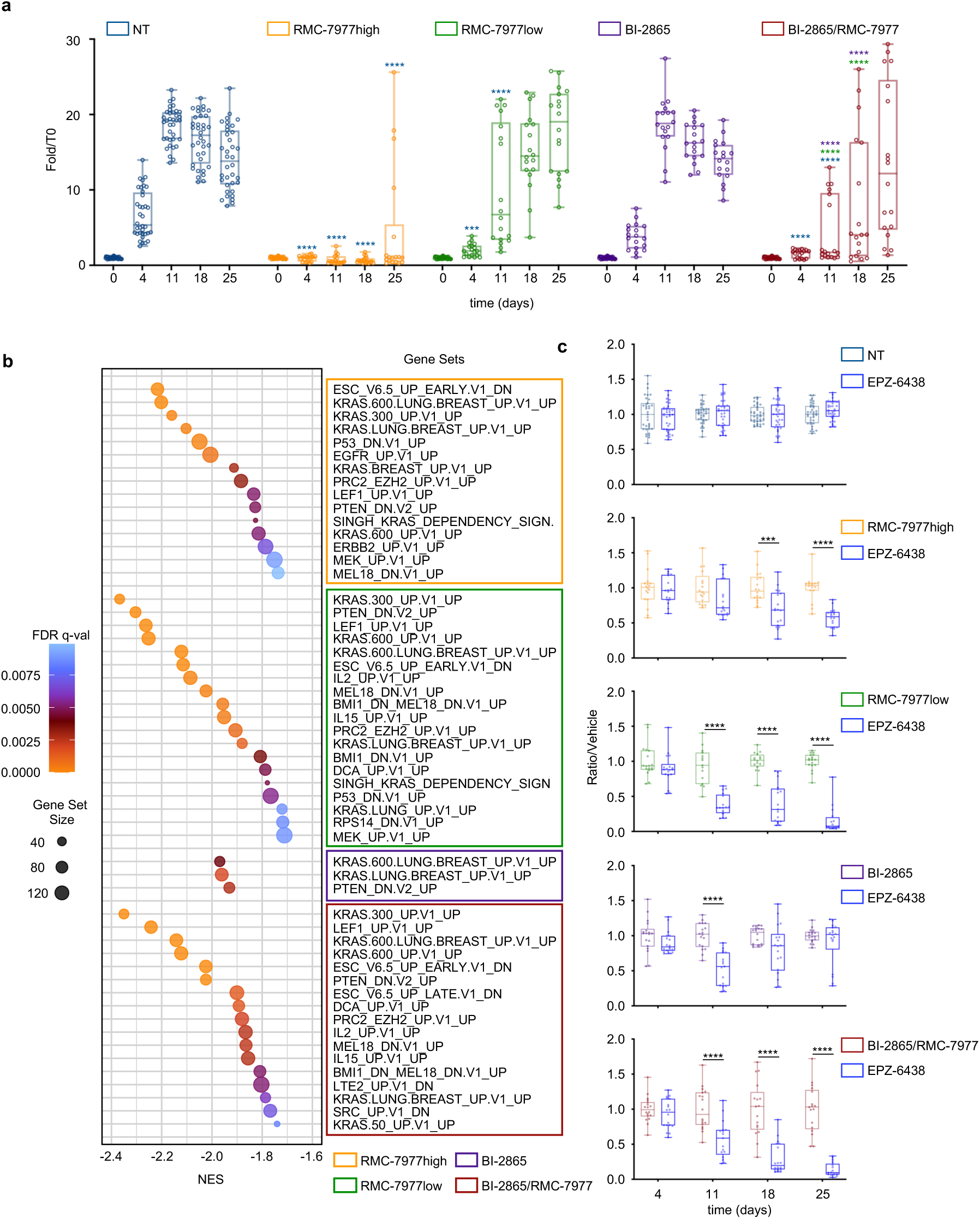
Modelling development of drug resistance in the MPO. **a**. The *KRAS*^G12A^-mutant PDO#17 was left untreated (NT) or exposed for 25 days to BI-2865 (10 μM), two concentrations of RMC-7977 (high: 5 μM and low: 0.5 μM) and a combination of BI-2865 and RMC-7977low (BI-2865/RMC-7977). At each time point, cell viability was evaluated by CellTiter-Glo 3D assay and reported as a fold change relative to the mean luminescence value at seeding time (T0). Each dot refers to the measurement performed in one single pillar (n= 18 for each treatment, n= 36 for NT). For each dataset, boxes show top and bottom quartiles, centre line denotes the median value, and whiskers indicate minimum to maximum. Results of 3 independent experiments are reported. A two-way ANOVA followed by Sidak’s multiple comparisons test was used to determine statistical significance between groups. Dark blue asterisks refer to the comparisons between the treatments and NT at the corresponding time point; in BI-2865/RMC-7977, green and purple asterisks refer to the comparisons with RMC-7977low and Bi-2865, respectively, at the corresponding time point. ***P= 0.0002; ****P< 0.0001. **b**. Organoids treated as in (a) were collected after 25 days of treatment. G&T-seq protocol was applied to obtain the paired separation and sequencing of DNA and mRNA of organoids recovered from pillars. Three independent pillars for each treatment were used. Dot plot depicts the enriched genes sets associated with KRAS inhibition as obtained by GSEA after sequence analysis of RNA. **c**. PDO#17 was treated as in (a) in presence or absence of EPZ-6438. Cell viability was evaluated by CellTiter-Glo 3D assay. For each combination and time point, luminescence values were normalised to the mean luminescence values of the corresponding vehicle-treated samples and presented as ratio/vehicle. Each dot refers to the data recovered from one single pillar (NT±EPZ-6438, n= 33-36; RMC-7977high±EPZ-6438, n= 16-18; RMC-7977low±EPZ-6438, n= 15-18; BI-2865±EPZ-6438 and BI-2865/RMC-7977±EPZ-6438, n= 17-18). For each dataset, boxes show top and bottom quartiles, centre line denotes the median value, and whiskers indicate minimum to maximum. Data from three independent experiments are reported. A two-way ANOVA followed by Sidak’s multiple comparisons test was used to determine statistical significance between groups. ***P= 0.0004; ****P< 0.0001.

The mechanisms of resistance to RAS inhibitors are still incompletely understood^70^. Hence, to gain insight on potential resistance mechanisms, we leveraged the G&T-seq protocol on organoids recovered at day 25th for all the treatment options. Genomic analyses demonstrated the overall similar mutational and copy number profiles between KRASi-treated organoids when compared with untreated samples (Supplementary Fig. 10a, c, d). Conversely, RNA-seq analysis revealed a profound reshape of the transcriptomic landscape following treatments (Fig. 5b). Specifically, gene-set enrichment analysis (GSEA) showed that upon prolonged drug exposure several cancer-relevant pathways were mostly downregulated, notably of RAS-centered networks, suggesting that RAS pathway inhibition was maintained in the resistant population, confirming previous results^71,72^. Intriguingly, we also observed down-regulation of hallmark targets of the Polycomb Repressive Complexes (PRCs) PRC1 and PRC2^73^ (Fig. 5b, Supplementary Fig. 10b and Supplementary Table 2). Since the histone methyltransferase EZH2, the catalytic component of PRC2, silences transcription at specific genomic sites by tri-methylating histone 3 at lysine 27 (H3K27me3)^74^, we hypothesised that PRC2-EZH2 activity might be involved in acquired resistance to RAS pathway inhibitors. To test this hypothesis, we repeated the experiment combining the EZH2 inhibitor EP-6438 (tazemetostat)^75^ alongside RAS inhibitors. Tazemetostat effectively reduced H3K27me3 deposition in the treated organoids (Supplementary Fig. 11a), albeit EZH2 inhibition by itself had no effect on cell viability even after prolonged exposure with high drug concentrations (10 μM) (Supplementary Fig. 11b). Conversely, combined treatment with RAS inhibitors resulted in a powerful synergistic effect on cell viability, with tazemetostat broadly enhancing the response to RAS pathway inhibition in all the tested conditions (Fig. 5c).

Collectively, these data support the use of the MPO platform for modelling resistance development, and reveal the potential of inhibiting EZH2 to overcome resistance to KRASi-based regimens.

### Proteomics, metabolomics, and lipidomics analyses on MPO-PDOs

A mechanistic understanding of the alterations triggered by treatment increasingly relies on integrated analyses encompassing not only genomic or transcriptomic profiling, but also proteomics, lipidomics and metabolomics alterations^76–78^. However, the routine adoption of such integrative multi-omics approaches is constrained by the substantial material requirements inherent to mass spectrometry (MS)-based platforms. This limitation is particularly critical in lipidomics and metabolomics, where analytical sensitivity and reproducibility are strongly dependent on input volume, extraction efficiency, and matrix complexity^79^. We hence explored whether the few microliters of PDOs present in each NEST pillar could yield enough material to support high-content MS-based multi-omics analyses.

An initial technical feasibility assessment was performed by comparing the results obtained by PDOs recovered from single domes cultivated conventionally, with those of MPO-grown organoids. We were able to retrieve from each MPO pillar, on average, 8,560 proteins, 229 lipids, and 33 polar metabolites, with an overlap of 73.26% and 52.40% with the bulk results, respectively for proteomics and lipidomics analysis (Supplementary Fig. 12a and 14a). Lower value (23.08%) was obtained for metabolomics (Supplementary Fig. 15a). Pairwise correlation plots (Supplementary Fig. 12b, 14b and 15b) further confirmed the biological and technical reproducibility across each pillar for these omics approaches.

We next explored whether these MS-multi-omics approaches might provide insights on the response of PDOs to treatment. Principal component analysis (PCA) on the proteomic profiles revealed a clear separation between the proteomes of non-treated versus FOLFIRI-treated organoids along the principal components (PCs) 2 and 3, confirming that proteomics is able to capture treatment impact (Fig. 6a). Notably, this separation was evident in both PDOs grown conventionally and in the MPO (Fig. 6a). Interestingly, the organoids recovered from pillars and the ones recovered from conventional domes separated along PC1, likely reflecting the expected variance due to differences in sample input and preparation format (Supplementary Fig. 12c). We next explored the pathways deregulated by the treatment, as defined by the proteome analysis. Importantly, we found overlapping pathways deregulated by the treatment in the two settings (Fig. 6b and Supplementary Fig. 12d, e), suggesting that the MPO, despite the reduced number of cells, nonetheless is able to capture the same proteomic alterations detected by conventional approaches bestowed with much higher sample availability. Intriguingly, the pathways emerging from the proteomic analysis were for the most part related to initiation of mRNA translation that were not captured by RNA analysis (Fig. 6c, Supplementary Fig. 13 and Supplementary Table 3), further highlighting the relevance of combining transcriptomics and proteomics analyses to comprehensively characterize cellular responses to drugs.

**Figure 6.**
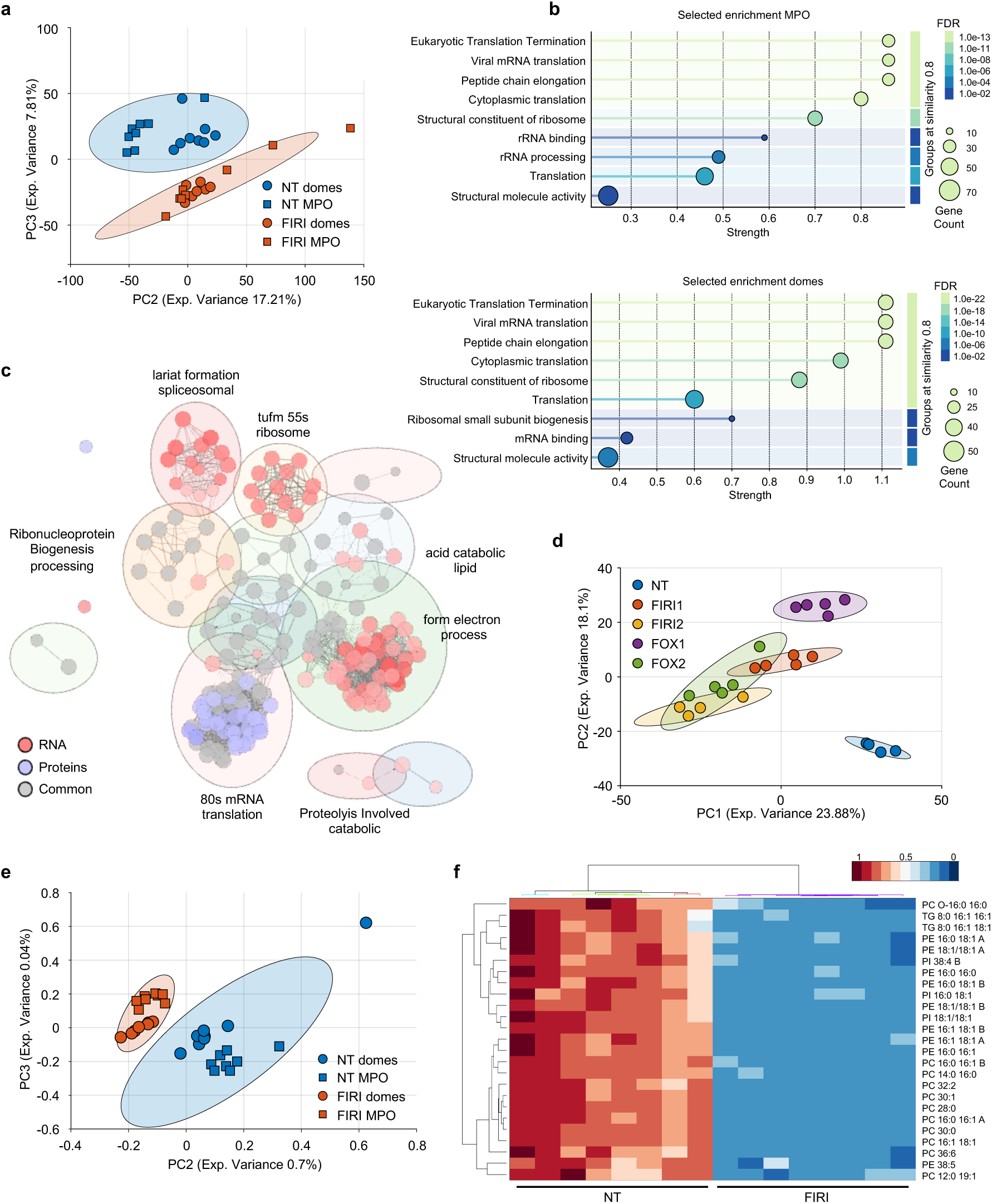
MS-multi omics analysis on organoids grown and treated in the MPO. a-c. PDO#61 was seeded in the MPO and left untreated (NT) or treated with FOLFIRI (FIRI) either in the MPO or in conventional conditions (domes) (n=8 for each experimental setting). At the end of the treatment, organoids were recovered and prepared for MS-multi omics analysis. (a) Score plot (PC2 *vs* PC3) from PCA of the proteomics datasets reporting the clustering of NT and FOLFIRI-treated organoids in the two experimental settings. (b) Reactome pathway enrichment analysis performed on statistically significant FOLFIRI-regulated proteins obtained from organoids grown in the MPO or in conventional conditions. (c) Network visualization of functional enrichment analysis performed on differentially expressed genes (adj. p-value <0.01) and proteins (adj. p-value <0.05), independently from fold change. Labels of the selected clusters indicate representative or common terms summarizing the biological processes enriched within each group, as determined by AutoAnnotate. **d**. PDO#61 was seeded in the MPO and left untreated or exposed to FOLFOX (FOX) and FOLFIRI (FIRI), used at two concentrations, one highest (FOX2 and FIRI2) and one 10–fold lower (FOX1 and FIRI1). Score plot (PC1 *vs* PC2) from PCA of the proteomics datasets reporting the clustering of NT and chemotreated-treated organoids. **e-f**. PDO#61 was seeded and treated as in a-c. (e) Score plot (PC2 *vs* PC3) from PCA of the lipidomics datasets reporting the clustering of NT and FOLFIRI-treated organoids in the two experimental settings. (f) Heatmap depicting the results of lipidomic analysis performed in NT and FOLFIRI-treated organoids recovered from the MPO. The variables shown were selected based on significance criteria from the corresponding volcano plot (Supplementary Fig. 14d, e).

Encouraged by these results, we expanded both the tested drugs and the concentrations used. Beside a clear distinction between treated and non-treated samples along PC1, there was a neat separation between the treatments with FOLFOX and FOLFIRI on PC2, albeit only at lower doses. Samples treated at high dosage clustered closer together, suggesting that at lower dosage the differences in the proteomic profiles reflect the different mechanisms of action of the two compounds, while at higher doses both compounds tend to converge on stereotyped apoptotic responses (Fig. 6d).

Also in untargeted lipidomics analysis, the PDOs recovered by MPO or grown in conventional domes were accurately separated after treatment with FOLFIRI, along the PC2 (Fig. 6e). As for the proteomics analysis, PC1 separated the PDOs grown by the two modalities (Supplementary Fig. 14c). Differential lipidomics analysis revealed that FOLFIRI treatment impacted glycerophospholipid remodeling, with a consistent decrease of phosphatidylcholines (PC) and phosphatidylethanolamines (PE) observed in both settings (Fig. 6f and Supplementary Fig. 14d-f), highlighting the potential role of lipid modulation in the response to chemotherapy^80^.

The assessment of the metabolome was less straightforward, as there was a less clear separation between untreated and treated PDOs within each platform (Supplementary Fig. 15c, d). Several metabolites including NAD+, Acetyl-L-Carnitine, Creatine and Guanidinacetic acid, showed the same trend across the two settings following FOLFIRI treatment (Supplementary Fig. 15 e-g), suggesting that the MPO is able to reliably capture metabolite alterations as more conventional approaches.

Overall, these results demonstrate that the MPO, combining drug sensitivity assays performed on PDOs with proteomics, lipidomics and potentially metabolomics studies, allows for better ex vivo modelling of drug responses. This underscores the potential of the MPO to more effectively uncover individual tumour vulnerabilities and predict tumour response to therapeutic agents.

## Discussion

We have designed an innovative microfluidic platform that allows the high-throughput screening of PDOs, overcoming several of the limitations that insofar have hampered the introduction of the PDOs technology both in the clinical practice and in drug discovery. While the standard practice for 3D cell culture privileges lab handwork and 48 or 96-plate format culturing^81–83^, we validated a system that combines 3D printing, microfluidics, and biological lab standards to build a fast, affordable, versatile, and robust drug screening platform on PDOs on a 384-plate format.

Some key features of the MPO include the seeding of organoids already embedded in ECM, the use of 3D printed biocompatible material for 3D culture supports and tools, and the straightforward integration of MPO into laboratory standards. This approach hence greatly simplifies the execution of survival, phenotypic and omic screenings, as well as of high-content imaging. Specifically, the NESTs culture setting, allowing a swift replacement of the culture medium and retrieval of the PDOs, overcomes limitations that restrict the scalability towards HTS by most of the emerging devices in the microfluidic and organ-on-a-chip field, as well as it addresses the growing need for standardisation of these technologies^84^, an important step towards the introduction of PDOs into the clinical practice.

Our approach is well suited to the “try-it-all” era of FPM, which holds the promise to deliver more effective and less toxic treatments for cancer patients^1,85^. Indeed, alongside several ongoing studies based on FPM^1,10^, a few functional clinical trials have already been implemented with remarkable successes^1,86^, even obtaining exceptional responses in heavily pretreated patients (EXALT trial^87^). Other studies are ushering new enticing paths, integrating ex-vivo screenings alongside extensive, in depth, multi-omic tumor profiling and clinical data, to predict patient responses^88,89^.

Identifying the most effective combination therapy in the clinical setting is challenging. It has been proven, in a large study involving over 1,000 PDX, that fewer than 5% of combinations improved progression-free survival more effectively than the best monotherapy^90^. In fact, much of the success of combination therapies^91^ has been attributed to the “bet-hedging” paradigm of independent drug action^92^, which posits that a single drug within a combination is primarily responsible for cancer cell killing in each specific patient. Therefore, MPO is among the tools that might assist in identifying the effective compound within a combination, sparing unnecessary toxicities to patients arising from the other, ineffective, drugs present in the combinations.

Beside bet-hedging, synthetic lethal drug combinations has also entered the clinic^93^, as for example the use of PARP inhibitors in BRCA-mutated cancers, the combination of BRAF and EGFR inhibitors in *BRAF*-mutant colon cancer and collateral lethality through co-deletions^94^ or co-amplifications^95^. While many promising combinations have been identified *in vitro*, their clinical translation has been hampered by two primary factors: reliance on inadequately predictive preclinical models^17^ – and the dependency on overly sensitive cell lines that inflate the perceived impact of additivity or synergistic interactions^92^. We argue that platforms such the MPO could overcome these limitations and finally lead to the preemptive identification of optimal single, or combination therapies.

The introduction of targeted therapies has heralded a new era in cancer treatment, but also posed new challenges. Unlike cytotoxic therapies, which rely on the maximum tolerated dose (MTD) as clinical guide, targeted therapies often do not exhibit dose-limiting toxicities. Thus, the MTD is often difficult to attain and less useful for determining clinical dosing^96^. Various strategies have been implemented to guide dosing, including target-mediated drug disposition (TMDD), and measurements of downstream effects^97^. In all, these methods rely on approximations often resulting in incomplete target engagement and suboptimal responses^92^. As such, undertreatment is very frequent, but at times also overtreatment, leading to toxicity-driven early discontinuation, where lower doses could instead be highly effective^92^. By integrating functional assays with multi-omic profiling and target engagement studies, even in a target-agnostic context, MPO delivers a rich molecular and functional landscape of drug response. This enables precise definition of optimal treatment concentrations and facilitates the discovery of next-generation therapeutic targets, a critical need for cancers lacking tractable oncogenes or clinically actionable biomarkers.

The ability of MPO to deeply model resistance development, uncovering the genetic and non-genetic mechanisms that tumor cells deploy to withstand therapeutic stress, offers powerful means to anticipate and overcome unexpected dependencies. We have effectively leveraged MPO to dissect the emergence of resistance to KRAS inhibitors, compounds that offer a transformative therapeutic opportunity across multiple cancers, yet remain plagued by the issue of resistance^70^. We identified a novel resistance mechanism driven by sustained PRC complex engagement. Pharmacological inhibition of its core enzymatic activity with Tazemetostat effectively counteracted the emergence of resistant clones. These findings establish a compelling rationale for epigenetic-based therapies, where combining KRAS and EZH2 inhibitors may offer a promising strategy to prevent resistance and improve outcomes in patients with KRAS-mutated mCRC.

Obtaining reliable images for phenotypic screenings from 3D models like PDOs has traditionally been challenging^47^. By adapting existing protocols, we enabled high-throughput implementation of these approaches including Cell Painting, on PDOs. These tools are increasingly integrated into drug discovery pipelines to predict compound activity, uncover novel therapeutic candidates and reveal hidden cancer vulnerabilities^25,55^. Importantly, within the MPO framework, these approaches require minimal sample processing and no prior knowledge of proteins and targets.

Cancer proteogenomics offers powerful insights into tumor biology and treatment response^98^, but its application at miniaturized scale requires innovative strategies to integrate functional and multi-omic data. This is particularly stringent with (MS)-based integrative multi-omics approaches constrained by the substantial material requirements. Through protocol miniaturization^63,99^, we demonstrated that MPO enables comprehensive pharmaco-proteogenomic analyses on the same sample. Despite the reduced input, MPO-grown organoids yield proteomics, lipidomics, and to a lesser degree, metabolomics profiles comparable to conventional systems, with omics data reliably distinguishing drug type, dose, and cellular impact. Future integration of more sensitive front-end separation techniques—such as nanoflow LC-HRMS, nano-LC or targeted approaches —will further enhance resolution and expand the utility of these approaches^100^.

We envision that the multi-layered, high-throughput analyses ushered by microfluidic platforms such as the MPO will have a profound impact both in the clinical practice, by defining drug sensitivities of patient-derived cells, in a time frame that could inform treatment decisions, as well as in large drug discovery programs to define the mechanisms of action of novel compounds.

## Acknowledgements

The authors wish to thank all the members of the Tonon, Dubini, Campiglia, Mazza, Zambrano, Tacchetti, Caravagna and Cittaro labs for helpful discussions and suggestions, alongside the clinical colleagues of the Aldrighetti team; Andrea Sottoriva and Nicola Valeri for helpful discussions, suggestions and for sharing reagents; Cesare Covino and Valeria Berno at the Advanced Light and Electron Microscopy BioImaging Center (ALEMBIC) core facility of IRCCS San Raffaele Scientific Institute and the San Raffaele Center for Omics Sciences (COSR) for expertise and assisting with instrumentation; Cinzia Felicita Sala for the generation of the Cohort Genomic Platform (C.G.P.). Supplementary Fig.1a and Supplementary Fig.5a were created at https://BioRender.com.

## Author contributions

O.A.B. designed and performed experiments, analyzed and interpreted data, coordinated the work and wrote the manuscript. E.B. designed and adapted the microfluidic tools and performed some experiments. J.M.B., C.F., G.F.G. performed experiments and analyzed data. E.S., F.M., V.C., V.G., D.L.G. and A.M. designed, performed and analyzed MS-multi-omics experiments. P.D.S. contributed to the design of the microfluidic tools. G.G, G.M. and G.Ga provided technical support to some biological experiments. V.G. performed bioinformatic analyses. V.R. and D.L. provided technical support to genomic experiments and analyzed data. E.C., P.F., S.Z., D.M. provided support to the imaging experiments and analyzed data. D.C. supervised bioinformatic analyses. S.B., managed patients’ data and participated in data discussion. F.P. performed pathology analyses on patients’ tissue samples and participated in data discussion. G.V.A. and G.C. performed some WES analyses. F.R. and L.A. collected surgical specimens and participated in data discussion. C.T. coordinated the work on imaging experiments and provided intellectual input. R.D.K. provided intellectual input. P.C. coordinated the work on MS-multi-omics experiments and provided intellectual input. G.D. coordinated the technical work on microfluidics and provided intellectual input. G.T. coordinated the work, provided intellectual input, interpreted data, and wrote the manuscript. All the authors revisited and accepted the final version of the manuscript.

## Material and methods

### The <u>M</u>icrofluidic <u>P</u>latform for <u>O</u>rganoids culture, MPO

The MPO consists of two main parts^32^: the “BMdrop dispenser”, includes a pump and a custom-made cartridge to deliver PDOs embedded in matrices directly into the 6x6 NESTs grids. Organoids are then placed in culture in a 384-well plate (Corning® Flat Clear Bottom White Polystyrene), each NEST in a well filled with 70 μL PDOs medium. The cartridge consists of a polymeric tubing specifically arranged in a cooled shell, to prevent the gelation and the sedimentation of the suspension of organoids in the biological matrices. The NEST is a 3D printed structure (Formlab, Form 3B+, Biomed Resin, SLA Stereolithography), designed to accommodate 8 µL of the sample inside a pseudo-cylindrical cavity, defined by eight pins and open top. In the culture configuration the NEST is hung, overturned and immersed in the 384-well plate, the 3D culture sample in the NEST is exposed to the cell culture media through the open side and the spaces between pins. The cartridge is connected to a syringe pump (PhD2000 Harvard Apparatus), which dispenses single drops of homogeneous PDOs suspensions into individual NESTs quickly and reliably, as demonstrated in our previous work^32^. Cartridges are individually packaged, sterilized, and autoclaved. Each set of NESTs is washed three times in deionized water, air dried at room temperature, and then UV sterilized under a biological hood.

### Human tissue samples

All procedures were approved by the Institutional Review Board of IRCCS San Raffaele Scientific Institute and were compliant with all relevant ethical regulations. Patient samples were collected at the Clinical Department in San Raffaele Hospital (Milan, Italy) under written informed consent in agreement with the Declaration of Helsinki (Protocols “ACC_ORG”, Milan, Italy). All data were stored according to best practices, protecting patient confidentiality and data integrity.

### Patient-derived mCRC organoids generation

Patient-derived organoids (PDOs) were obtained essentially as already reported^101^. Briefly, surgical resections of LM-CRC were minced, and digested in 5mM PBS/EDTA supplemented with 2X TrypLe (Gibco, A1217701) and 500U recombinant DNase I (Roche, 04716728001) at 37°C for 60 min. Cells released in the solution were resuspended in 90 μL growth factor reduced Matrigel^®^ (Corning, 356231) and seeded in a 24-well flat bottom cell culture plate (Corning). After solidification, Matrigel^®^ domes were overlaid with Advanced DMEM/F12 (Gibco, 12634028) supplemented with 1X B27 (Gibco,17504044), 1X N2 (Gibco, 17502048), 1% penicillin-streptomycin (Euroclone, ECB3001D), 1% glutamine (Gibco, 25030081), 1% BSA (Sigma-Aldrich, 10735086001) and with the following growth factors: 50 ng/mL EGF (PeproTech, 100-39H), 100 ng/mL Noggin (PeproTech, 250-38), 500 ng/mL R-Spondin 1 (PeproTech, 120-38), 10 nM gastrin (Sigma-Aldrich, G9145), 10 ng/mL FGF-10 (PeproTech, 100-26), 10 ng/mL FGF-basic (PeproTech, 100-18B), 100 ng/mL Wnt-3A (R&D Systems, 5036-WN), 1 μM prostaglandin E2 (Tocris Bioscience, 2296), 10 μM Y-27632 (StemCell Technologies, 72304), 4 mM nicotinamide (Sigma-Aldrich, N0636), 0,5 μM A83-01 (Tocris Bioscience, 2939), 5 μM SB202190 (Sigma-Aldrich, S7067). Medium was changed every 2–3 days.

### Seeding and culturing of PDOs in the MPO

Sub-confluent PDOs were harvested, mechanically and enzymatically dissociated into single cells by incubating with TrypLE select enzyme and replated^101^. Cells were allowed to generate small PDOs for 2-3 days and then recovered with the Cell Recovery solution (Corning, 354253). Recovered PDOs were diluted in a solution composed of ∼5 mg/ml Matrigel^®^ or Cultrex Basement Membrane Extract (BME, R&D Systems, 3432-010-01) in PBS. The suspensions were set to obtain ∼500-3.000 cells aggregated in organoids within each 8 µL NEST unit. The suspension of PDOs was then loaded into the BMdrop dispenser. Organoids were seeded onto the NESTs grids and cultured as described above. For long-term culture, PDOs diluted in ∼7 mg/ml Matrigel^®^ or BME were plated in the MPO and maintained in culture for up to 25 days. Medium was refreshed twice a week by placing NESTs grids in a new 384-well plate with fresh medium. Growth of organoids was evaluated at fixed points analysing organoid viability by CellTiter-Glo 3D assay (see below) and brightfield imaging analysis (Zeiss Axio Observer Z1). The culture was stopped when no further increase in the luminescence values was observed, indicating that the culture had reached a balance between cell viability and death.

### Drug screening and cell viability assay

Oxaliplatin and 5-FU were obtained from the pharmacy at San Raffaele Hospital and dissolved in PBS. SN-38, the active metabolite of irinotecan (Sellekchem, S4908), was dissolved in DMSO (Sigma-Aldrich, D8418). Compounds were kept at stock concentrations of 1000X in the vehicle. Before treatments, drugs were further diluted to 1X final concentration in complete PDOs medium and 70 μL of this solution were added in each well. DMSO or PBS treated PDOs served as control. DMSO concentration in the medium was 0.1%. For the retrospective and prospective studies PDOs were seeded in the platform and exposed to a combination of drugs that replicated two chemotherapeutic regimens commonly used to treat patients with mCRC, FOLFOX (FOX) and FOLFIRI (FIRI). Two concentrations were used, one highest (FOX2 and FIRI2) based on the Cmax reported for each drug^37–40^ and one 10–fold lower (FOX1 and FIRI1), as follow:

FOX1: 40 μM 5-FU and 1.25 μM Oxaliplatin

FOX2: 400 μM 5-FU and 12.5 μM Oxaliplatin

FIRI1: 40 μM 5-FU and 10 nM SN-38

FIRI2: 400 μM 5-FU and 100 nM SN-38

One organoid line (PDO#119) was exposed to treatments replicating the FOLFOXIRI combination, at two concentrations:

T1: 40 μM 5-FU, 1.25 μM Oxaliplatin and 10 nM SN-38

T2: 400 μM 5-FU, 12.5 μM Oxaliplatin and 100 nM SN-38

Organoids were incubated with the compounds at 37°C 5% CO2 for 96 hours with no further changes of media or compounds refresh. After treatment PDOs were displaced from the chip into a new empty 384-well white plate (Corning), by centrifugation (3 minutes at 800 x g) and cell viability was quantified by CellTiter-Glo 3D assay (Promega, G9681) essentially following the manufacturer’s instructions. Briefly, a 1:1 mix of CellTiter-Glo 3D and Advanced DMEM/F12 was added to organoids and incubated for 45 min at RT. The luminescence signal was then measured on a Mithras LB 940 microplate reader (Berthold Technologies). Mean of the luminescence values of the six technical replicates was calculated for each condition and reported as percentage (± standard deviation [s.d.]) of viable cells relative to the untreated samples. Then, mean ± Standard Error of the Mean (s.e.m.) between biological independent experiments was calculated. Drug screenings were repeated at least twice and treatments performed in six technical replicates. In the prospective study, the optimal time for drug screening was defined based on the organoids growth rate and size (50-100 µm)^102^.

### Homogeneous time resolved fluorescence (HTRF)

The KRAS inhibitor BI-2865 (#HY-153724) was purchased by MedChemExpress and dissolved at a 1000X concentration in DMSO. Detection of ERK expression and phosphorylation levels was performed using the total ERK (64NRKPEG) and the phospho-ERK (Thr202/Tyr204) (64ERKPEG) cellular kits from Revvity. Organoids were seeded and grown in complete medium for 4 days. Before treatment the chip containing PDOs was placed in a 384-well plate with a minimal medium where growth factors were withdrawn, for 20 hours. After this time, the chip was placed in a new plate containing fresh minimal medium in which either DMSO or BI-2865 at the final concentration of 10 μM were already dissolved. Treatment was performed for 4 hours, then the chip was centrifuged on a 384-well plate (3 minutes at 400 x g) and the assay performed essentially following the manufacturer’s instructions. 16 μL of 1X lysis Buffer (including 1X blocking reagent) was added to each well for 45 minutes at room temperature. After lysis, 4 μL of a 1:1 mixture of d2 and Eu Cryptate antibodies working solution was added in each well. Plates were sealed and incubated overnight at room temperature light protected. The HTRF program was used to detect signals at 620 nm and 665 nm wavelengths using a Mithras LB 940 microplate reader (Berthold Technologies, Bad Wildbad, Germany). Negative control to check the non-specific signal and positive control as provided by the manufacturer, were used in each assay.

### Immunoblots and antibodies

Whole-cell extracts were obtained by lysis in sodium dodecyl sulphate (SDS) buffer (50 mM of Tris HCl pH 6.8, 10% glycerol, 2% SDS). Proteins were separated by SDS-polyacrylamide gel electrophoresis, blotted onto an Nitrocellulose membrane (Merck KGaA, Darmstadt, Germany), and probed with the indicated antibodies. Clarity Western ECL Blotting Substrate (Bio-Rad, Hercules, CA, USA) was used for the chemiluminescent reaction. The antibodies used were as follow: Phospho-p44/42 MAPK (Erk1/2) (Thr202/Tyr204) (#9101), p44/42 MAPK (Erk1/2) (#9107) and Tri-Methyl-Histone H3 (Lys27) (#9733) from Cell Signaling (Leiden, The Netherlands); anti-Vinculin (V9131) from Sigma-Aldrich; HRP-conjugated anti-mouse and anti-rabbit secondary antibodies from GE Healthcare. ImageLab software (ver 6.1) was used for image analysis and quantitation.

### PDOs imaging and data analysis

For immunofluorescence staining, PDOs were plated on NESTs grid supports specifically designed for imaging applications and exposed to drugs as described above (FIRI2 for 96 hours). After treatment, PDOs were stained directly into the NESTs grid. All steps were performed at room temperature and solutions were prepared in PBS. PDOs were fixed in a solution of 4% paraformaldehyde (Electron Microscopy Sciences, 157-8) and 2% sucrose (Sigma-Aldrich, S0389) for 30 minutes and then permeabilized with 0.1% Triton X-100 (Sigma-Aldrich, T8787) for 8 minutes. Nonspecific antigens were blocked in a solution of 5% BSA (Sigma-Aldrich, A8022) and 10% goat serum (Sigma-Aldrich, G9023) for 30 minutes, and then PDOs were incubated with monoclonal antibody γH2AX (Biolegend, 613402) at 1:100 dilution in 2.5% BSA for 4 hours. After incubation for 2 hours with a solution of goat anti-mouse IgG Alexa Fluor™ 488 (Invitrogen, A-11001) at 1:500, 10μg/mL Hoechst 33342 (Invitrogen, H3570) and 4μg/mL rhodamine phalloidin (Invitrogen, R415) in 2.5% BSA, the NESTs grid was placed in a new empty 384-well black glass bottom microplate (Ibidi, 88407) and PDOs were centrifuged at 400 x g for 3 minutes to collect the individual drops within the wells. Drops were overlaid with PBS, the microplate sealed with an optical adhesive film (TermoFisher, 4311971) and stored at 4°C prior to imaging. 16 bit −1024x1024 pixels images were acquired using a 60X silicone immersion objective on an Olympus Fluoview 3000RS (Olympus Life Sciences) confocal microscope. Nuclei were manually segmented in ImageJ (The Mathworks Inc.) to estimate nuclear area and the average fluorescence intensity per nucleus in the γH2AX channel was evaluated.

### Spheroids culture live imaging and data analysis

MCF7 stably expressing p65-mScarlet construct, were generated by lentiviral infection. Federica Colombo and Alessandra Agresti kindly shared plasmids and protocol for p65-mScarlet lentiviral infection and HEK-293T cells were used for virus production. Infected MCF7 cells were sorted for high mScarlet signal and NF-kB activation upon TNFα was validated via confocal imaging following the previously described procedure for live cell imaging in 2D^51^. To allow spheroid formation, mScarlet-p65 MCF7 single cells were plated on uncoated low-attachment dishes with a density of 300 cells/cm^2^ and cultured for 5 days. Before imaging, spheroids were embedded in Matrigel and plated in the NESTs, treated with 20 ng/ml TNF or vehicle control and live imaged using Olympus Fluoview 3000RS confocal microscope with temperature and CO_2_ control. Quantification of the evolution in time of nuclear to cytosolic intensity of p65 was performed by adapting a procedure already described^51^. First, nuclear and cytosolic masks of a plane of the 3D stack images were identified using CellPose^103^. Using the baricenters of the nuclei we tracked the cells using an adaptation of the Hungarian algorithm and then we calculated the background-subtracted nuclear to cytosolic average intensities for each track cell based on the nuclear and cytosolic masks. This gives values well below 1 for untreated cells (NT) and close to 1 for TNFα-treated cells, as previously reported^51^. Area under the curve (AUC) was calculated for single nuclear to cytosolic p65 profiles of 2 hours duration.

### Phenotypic profiling by Cell Painting assay

Organoids and U2OS cells were treated with: CA-074 methyl ester 10 μM, Tetrandrine 10 μM, Fenbendazole 10 μM, Berberine sulfate 20 μM, Etoposide 10 μM and Cytochalasin B 20 μM all from MedChemExpress. Organoids were also treated: with BI-2865 10 μM and RMC-7977 (#HY-156498 from MedChemExpress) at 5 μM, FOX2 and FIRI2 (see above). U2OS cells were seeded at the density of 1000 cells per well into a 384-well microplate (Ibidi, 88407) incubated at 37 °C, 5% CO2 overnight and then treated for 48 hours. PDO#17 was plated in the MPO as described above and treated for 24 and 48 hours. Morphological profiling was performed using the Cell Painting JUMP kit (Revvity, PING21), for U2OS cells following the manufacturer’s instructions, for organoids by adapting a protocol previously reported^47^. Organoids were first incubated with PhenoVue 641 Mitochondrial Stain (500 nM) for 1 hour at 37°C and then fixed as described above. Subsequently, organoids were stained for PhenoVue Hoechst 33342 Nuclear Stain (1.62 μM), PhenoVue Fluor 488 - Concanavalin A (48 nM), PhenoVue 512 Nucleic Acid Stain (6 μM), PhenoVue Fluor 555 – WGA (43.7 nM), PhenoVue Fluor 568 – Phalloidin (8.25 nM), in a solution of 0.1% Triton X-100 in PBS for 2 hours at room temperature. Then organoids were displaced from the NESTs grid, collected onto a new empty 384-well black glass bottom microplate as individual drops within the wells and stored as described above prior to imaging. An Olympus Fluoview FV3000 confocal laser scanning microscope with a silicon-oil 30X/1.05NA objective was used to collect confocal z-stacks (pixel size = 0.207 μm/pixel, z-step = 1 mm) with sequential channels acquisition: laser 405 nm for Hoechst 33342, laser 488 nm for PhenoVue Fluor 488, laser 514 nm for PhenoVue 512, laser 561 nm for AlexaFluor 555, laser 594 nm for Alexa Fluor 594m and laser 640 nm for Cy5. Single-plane images at the focal plane of the maximum organoid section were collected for the follow-up analysis. Analysis was carried out by segmentation and feature extraction using a custom pipeline in CellProfiler^104^ that identifies the whole cell, cytoplasm and nucleus using the Phalloidin and the Hoechst images, and then computes the single-cell features for each identified compartment and for each staining. The resulting features are then aggregated into medians at the per-well level, normalized, and reduced as already described^54^. To avoid overfitting and reduce feature redundancy, we computed pairwise correlations between feature vectors and performed hierarchical clustering on the resulting distance matrix and cut the resulting tree at distance 1. For each resulting cluster we selected the feature with the highest mutual information with the treatment label as representative feature. Finally, the feature matrix was used to fit a Partial Least Squares regression (PLS) model with treatments as target label, using python scikit-learn PLSCanonical function (parameters: n_components = 5, scale = True).

### Whole Genome Sequencing of PDOs grown in conventional conditions

DNA was extracted using the AllPrep DNA/RNA Mini kit (Qiagen) following manufacturer’s instructions. Library preparation was conducted using a non-PCR-free protocol and performed by Novogene using Novogene NGS DNA Library Prep Set (Cat No.PT004). 150bp paired-end sequencing (60X coverage) was carried out by Novogene on Illumina platforms according to the effective library concentration and required data amount.

### Separation and parallel sequencing of the genome and transcriptome using the G&T-seq protocol

Genomic DNA and mRNA separation was performed according to G&T-seq protocol^105^. Organoids were seeded in the MPO and treated for 24 and 72 hours. At each time point, to minimize the DNA contamination of Matrigel, organoids were treated with RQ1 RNase-Free DNase (Promega, cat. no. M6101) for 30 minutes at 37°C directly by immersing the NESTs grids in the solution placed in the 348-well plate. After washes with PBS, the single Matrigel drops were collected by rapid centrifugation in 50 µL of RLT plus lysis buffer (Qiagen, 1053393). Lysed cells were transferred into 200ul tubes and incubated for 20 minutes with 10 µL of oligo-dT conjugated to streptavidin beads (Invitrogen, 65001). After incubation, the poly-adenylated mRNAs were separated using a magnet, while the DNA, present in the supernatant, was transferred to a new tube. Reverse Transcription (RT) was performed following manufacturer’s instructions. Next, the cDNA was amplified by directly adding 7.5 μL of PCR reaction mastermix (6,25 μL Kapa HiFi HotStart ReadyMix (2X) (Kapa, KK2601) and 0,12 μL IS PCR primers (5-AAGCAGTGGTATCAACGCAGAGT-3; IDT) with following thermal protocol: 98°C for 3 min; 16 cycles of 98°C for 20 sec, 67°C for 15 sec and 72°C for 6 min; 72°C for 5 min and hold at 4°C. Finally, the amplified cDNA and genomic DNA (gDNA) were purified with AMPure XP beads (1X ratio) (Beckman Coulter, A63881). Quality check was performed using Tape Station platform (Agilent Technologies) in combination with Qubit High sensitivity DNA Assay Kit (Agilent Technologies, 5067-4626). DNA and RNA libraries were prepared using Nextera XT kit (Illumina, FC-131-1096) according to manufacturer’s instructions. After library preparation, samples were purified using AMPure XP beads (1x ratio) and quality check was performed as described above. Library pools were sequenced on NovaSeq 6000 (Illumina) at low coverage (0.1X; paired-end, 100 bp) for DNA and at a depth of 30 million paired-end (2X100 bp) for RNA. Read tags were aligned to the hg38 reference genome using bwa^106^ (for DNA) and STAR^107^ (for RNA) with default parameters. Copy Number Alterations were identified in DNA alignments using CNVPytor^108^, using variants called with bcftools^109^ filtered for quality higher than 15.

### Whole exome sequencing

Library generation was performed using the Twist protocol according to the manufacturer’s guidelines (Twist Bioscience). Fifty nanograms of gDNA were enzymatically shattered, end-repaired and dA-tailing to generate dA-tailed DNA fragments. Twist universal adapters were subsequently added to generate gDNA libraries ready for index introduction via amplification with Twist UDI Primers. At each step, the products were purified. Quantification and sizing of each library were done using the Qubit dsDNA (Thermofisher) and D1000 Tape Station assay (Agilent Technologies). The exome DNA was enriched by Comprehensive Exome Enrichment (Twist Bioscience). After hybridization, the targeted molecules were captured on streptavidin beads and amplified. The resulting enriched DNA libraries were purified and quality control was assessed using D1000 high sensitivity Tape Station assay (Agilent Technologies). Pools were run on NovaSeq6000 (Illumina) at a depth of 40X paired-end (2X100 bp). For analysis on PDO#61 and #74, read tags were aligned to the hg38 reference genome using bwa^106^. Genotype discordance of WES and RNA-seq with high coverage WGS was evaluated using bcftools, setting the error probability to 0, hence allowing raw comparison of the mismatched variants by their likelihood.

### Modelling of drug resistance development in the MPO

PDO#17 was seeded within the MPO at a concentration of approximately 1000 cells/pillar. PDO#17 was then treated with BI-2865 10 μM, RMC-7977 5 μM (high), RMC-7977 0.5 μM (low) and EPZ-6438 10 μM (Tazemetostat, #HY-13803 from MedChemExpress). Compounds were kept at stock concentrations of 1000X in the vehicle, DMSO. Before treatments, drugs were further diluted to 1X final concentration in complete PDOs medium and 70 μL of this solution were added in each well of a 384-well plate. DMSO-treated PDOs served as control. Organoids were treated for 25 days. Medium was refreshed twice a week by placing NESTs grids in a new 384-well plate with fresh medium. Growth of organoids was evaluated at fixed points analysing organoid viability by CellTiter-Glo 3D assay (see above). On day 25th organoids were recovered and genomic DNA and mRNA separation, sequencing and bioinformatics analyses were performed according to the G&T-seq protocol (see above). Genomic DNA was also subjected to WES analysis. Library preparation was carried out as described above. Mutation calling was performed using the nf-core pipeline Sarek 3.5.1^32,110^ on the ORFEO cluster from Area Science Park. Variant calling was performed using Mutect2 in multisample mode^111^, while copy number calling was performed using ASCAT^112^ in accordance with the Sarek recommendations for running it on WES samples^113^. Variant annotation was performed by using the Ensembl-VEP^114^ with cache version 110 from the corresponding reference genome. Mutations and copy number data were processed with in-house tools and scripts. COADREAD driver list was downloaded by the IntOGen database to identify potential driver genes involved in the onset and evolution of the disease. Positions with less than 20 reads mapped in any of the samples had been removed from the analysis, as considered lowly informative.

### Gene set enrichment analysis (GSEA)

Unbiased gene set enrichment analysis was performed using GSEA software developed by Broad Institute^115,116^ on differential expressed genes pre-ranked by fold change with 2,000 permutations. Reference gene sets were obtained from the MsigDB library for C6 collection: oncogenic signature gene sets^117^. Gene sets with a FDR q-value < 0.01 were considered significant.

### Organoids preparation for MS-multi omics analysis

Organoids were seeded and treated in the MPO as described above for 72 hours. After treatment, organoids were displaced from the chip into a new empty 384-well solid plate with conical bottom (Greiner #781280) by centrifugation (3 minutes at 800 x g). Then 100 µL of Cell Recovery solution was added in each well. After 45 min at 4°C, organoid pellets were washed once with PBS, snap frozen and stored at −80°C. In a parallel experiment, organoids were grown and treated in conventional conditions, one dome being seeded with 16X more organoids than one pillar.

Each organoid sample, whether recovered from a conventional dome or a single pillar, was subjected to a simultaneous extraction of protein, polar metabolites and lipids according to a previous protocol with minor modifications^118^. To avoid interference from the thousands of identical peptides shared between biological matrices and organoids^119^, we performed cell recovery based washes. Subsequently, 226 µL of ice-cold methanol (MeOH) containing a blend of deuterated standards was spiked to organoid pellets kept at − 30 °C for 1 minute and then transferred to an ultrasonic bath for 10 minutes. Following the addition of 750 µL of ice-cold methyl tert-butyl ether (MTBE), cells were incubated in a Thermomixer (Eppendorf) for 1 hour at 4 °C and 550 rpm. 188 µL of water (H2O) was added to the samples, and after centrifugation, the upper (for lipids) and lower (for metabolites) layers were collected and evaporated using a SpeedVac (Savant, Thermo Scientific, Milan, Italy). Dried extracts were dissolved in 150 μL of CHCl3/MeOH/IPA (1/2/4) and 50 μL of H2O/MeOH 90:10 (v/v%), respectively, for lipidomics and metabolomics analyses. Remaining protein pellet digestion and clean-up was performed as reported previously^120^.

### Proteomics profiling

Proteomic analysis was performed by nLC- HRMS using an Ultimate 3000 nanoLC (Thermo Fisher Scientific, Bremen, Germany) coupled to an Orbitrap Lumos tribrid mass spectrometer (Thermo Fisher Scientific) with an Easy nano electrospray ion source (Thermo Fisher Scientific). Peptides were trapped for 1 minute in a PepMap trap-Cartridge, 100 Å, 5 µm, 0.3 x 5 mm (Thermo Fisher), and separated with a EasySpray PepMapC18 column (250 mm × 75 μm I.D, 2.0 µm, 120Å, IonOpticks, Fitzroy, Australia). Mobile phases were A): 0,1% HCOOH in water v/v; B): 0,1% HCOOH in ACN/Water v/v 80/20. Peptides were separated using a linear gradient of 90 min. HRMS analysis was performed in data dependent acquisition (DDA) as follows: DDAparameters: MS1 range 400–1500 m/z, HCD fragmentation was used with normalized collision energy setting 27. Resolution was set at 120.000 for MS1 and 15.000 for MS/MS. Single charge and unassigned charge peptides were excluded. Quadrupole isolation was set to 3Da. Maximum ion injection times for MS (OT) and the MS/MS (OT) scans were set to auto and 50 ms respectively, and ACG values were set to standard. Dynamic exclusion: 30 s. DDA raw MS data were analysed using Proteome Discoverer v 2.5 (Thermo Fisher), Proteins were considered identified with at least one unique peptide, using a false discovery rate (FDR) threshold of 1. The following parameters were used: and 0,6 Da for precursors and fragments respectively. Sequest search and Percolator algorithms were used. Carbamidomethylcysteine was used as fixed modification while methionine oxidation as variable. Proteomics enrichment was performed with STRING^121^.

### Lipidomics profiling

Lipidome has been profiled using a Thermo Ultimate UHPLC system (Thermo Scientific, Bremen, Germany) connected online to a TimsTOF Pro Quadrupole Time-of-Flight (Q-TOF) mass spectrometer (Bruker Daltonics, Bremen, Germany) equipped with an Apollo II electrospray ionization (ESI) source. Lipids separation was performed using a Waters Acquity UPLC CSH™ C18 column (100 × 2.1 mm; 1.7 μm, 130 Å) paired with a VanGuard CSH™ precolumn (5.0 × 2.1 mm; 1.7 μm, 130 Å) (Waters, Milford, MA, USA). The column oven temperature was maintained at 65 °C, with a flow rate of 0.55 mL min^-1^. The mobile phases were composed of: (A) ACN/H_2_O 60:40 (*v/v %*) and (B) IPA/ACN 90:10 (*v/v %*), both buffered with 10 mM HCOONH_4_ and 0.1% HCOOH (*v/v %*) for the positive ionization mode and with 10 mM CH_3_COONH_4_ and 0.1% CH_3_COOH (*v/v %*) for the negative ionization mode. The following gradient was applied: 0 min, 40% B; 0.4 min, 43% B; 0.425 min, 50% B; 0.9 min, 57% B; 2.0 min, 70% B; 2.950 min, 99% B; 3.3 min, 99% B; 3.31 min, 40% B, followed by a 0.7 min re-equilibration of the column. At the start of each run, mass and mobility accuracy were recalibrated by injecting a mixture (1:1 *v/v %*) of 10 mM sodium formate calibrant solution and ESI-L Low Concentration Tuning Mix. Data-dependent parallel accumulation serial fragmentation (DDA-PASEF) acquisition mode was employed. Both positive and negative ESI ionization were used in separate runs, with each sample analyzed in duplicate. The injection volume was set at 2 μL and 3 μL, respectively, for positive and negative ionization mode. ESI source parameters were set as follows: nebulizer gas (N_2_) pressure at 4.0 bar, dry gas (N_2_) flow rate at 10 L/min, and dry temperature at 280 °C. Mass spectra were collected over the m/z range of 50–1500, with accumulation and ramp times of 100 ms each. Ion mobility was scanned from 0.55 to 1.80 Vs/cm². Precursors were isolated within ± 2 m/z and fragmented using ion mobility-dependent collision energies MS/MS stepping mode: 1/K0 [Vs/cm²] 0.55-1.8; CE [eV] #1: 20–40 and CE [eV] #2: 35–50. The total acquisition cycle lasted 0.32 seconds, consisting of one PASEF ramp. Exclusion time was set to 0.1 minutes, and ion charge control (ICC) was configured to 7.5 million. Data alignment, filtering, and annotation were carried out using MetaboScape 2023b (Bruker) that uses a feature-detection algorithm (T-Rex 4D). The feature detection threshold was set to 250 counts for both positive and negative ionization modes, with a minimum of 75 data points in the 4D-TIMS space. Lipidomics data were deconvoluted in positive mode using [M+H]^+^, [M+Na]^+^, [M+K]^+^, [M+H–H_2_O]^+^ and [M+NH_4_] ^+^ ions, and in negative mode using [M−H] ^−^, [M+Cl] ^−^, [M+CH_3_COO]^−^ and [M−H_2_O] ^−^ ions. Lipids were annotated through a rule-based approach and the LipidBlast spectral library in MS-DIAL. Molecular formulas were assigned using SmartFormula^TM^ (SF). Compound annotation has been performed setting the following parameters: mass accuracy (narrow: 2 ppm, wide: 10 ppm), mSigma (narrow: 30, wide: 250), MS/MS score (narrow: 800, wide: 150), and Collision Cross-Section (CCS) % (narrow: 2, wide: 3.5). CCS values were compared with predictions from the CCSbase platform. After the annotation process, all spectra were manually curated and examined. Manual curation of each lipid was then performed following Lipidomics Standard Initiative (LSI) guidelines^122^, specifically, besides MS/MS diagnostic ions, crucial aspects of lipid annotation such as (a) lipid adducts in electrospray ionization and (b) regular retention behavior, e.g., the equivalent carbon number (ECN) model used for RPLC, were carefully evaluated. LipidCreator tool^123^ extension in Skyline^124^ was used for in silico comparison of specific product ions. Lipids missing in more than 75% of real samples and 50% of QC samples were excluded. Additionally, lipids with a Coefficient of Variation (CV) above 30% among QCs were discarded. Lipid intensities were normalized using the corresponding deuterated internal standard. The dataset was then log-transformed and autoscaled for multivariate data analysis. Functional lipidomics analysis was performed with LipidOne^125^.

### Metabolomics profiling

Metabolome analyses were performed on a Thermo Vanquish Neo UHPLC (2.1 mm I.D setup) coupled online to a hybrid quadrupole Orbitrap Exploris 120 mass spectrometer (Thermo Fisher Scientific, Bremen, Germany) equipped with a heated electrospray ionization probe (HESI II). The MS was daily calibrated by Pierce™ FlexMix™ calibration solutions in both polarities. Metabolites separation was performed with an Acquity UPLC HSS T3™ (150 × 1.0 mm; 1.8 µm, 100Å), the column temperature was set at 55 °C, a flow rate of 100 µL/min was used, mobile phase consisted of (A): H_2_O + 0.1% HCOOH and (B): ACN + 0.1% HCOOH were used for positive ionization, while 0.1% CH_3_COOH was used for negative ionization mode. The following gradient has been used: 0 min, 0% B; 6 min, 70% B; 8 min, 80% B; 9 min, 98% B, 10 min 98% B; 10.1 min 0% B and 3.9 min for column re-equilibration. The HESI source parameters for 1.0 mm i.d setup were: sheath gas 20 a.u, auxiliary gas 7 a.u, sweep gas 0. Spray voltages were set to 3.3 kV and 3.0 kV for ESI (+) and ESI (-) respectively, Ion transfer tube and vaporizer temperature were set to 280° and 150°C respectively, in both RP and HILIC modes. MS data acquisition for both setups was performed in full scan-data dependent acquisition (FS-DDA) in the m/z 70-800, MS1 resolution was set to 60000, the AGC target was set to auto with a maximum injection time at 100 ms. MS/MS was employed with an isolation window of 1-5 Da, dynamic exclusion of 10s, resolution was set to 15.000, and HCD was used with normalized collision energies of 20, 40 and 60. The instrument was externally calibrated daily with FlexMix solution (ThermoFisher) while at the beginning of every LC run the internal calibrant was injected (IC run start mode). FreeStyle (Thermo Fisher Scientific) was used to visualize RAW data, which were then imported to Compound Discoverer v.3.3 (Thermo Fisher Scientific) to normalize, align, detect and identify compounds. Features were extracted from 0-10 min and 0-11 min of the HILIC and RP chromatography runs, respectively, in the m/z= 70-800 mass range. Data were aligned according to an adaptive curve alignment model. Compounds were detected using the following parameters settings: mass tolerance was set to 5 ppm, while retention time tolerance was set to 0.2 min; minimum peak intensity was set to 100000 AU and the signal-to-noise threshold for compound detection was set to 5. The peak rating filter was set to 3. To perform blank subtraction, we maintained max sample/max blank ratio > 5. For statistical comparison between the two setup, signal intensity was normalized by using the algorithm “Constant Sum”. To predict elemental compositions of the compounds, the relative intensity tolerance was set to 30% for isotope pattern matching. For the mzCloud database search, both the precursor and fragment mass tolerance were set to 5 ppm. The databases used for matching compounds in ChemSpider for structural search were BioCyc, the Human Metabolome Database and KEGG, and the mass tolerance in ChemSpider Search was set to 5 ppm. The mass tolerance for matching compounds in Metabolika pathways was set to 5 ppm. Compounds were assigned by comparing annotations using the following nodes in order of priority: (1) mzCloud (2) Predicted Compositions; (3) MassList search; (4) ChemSpider Search; (5) Metabolika search.

### MS-multi-omics data analysis

Data analysis was performed using MATLAB (The MathWorks, Inc., Natick, MA, USA), combining standard and custom scripts developed for this study. The analytical pipeline included data preprocessing, statistical analysis (both univariate and multivariate), and visualization. To investigate the technical feasibility of the MPO approach, a comprehensive multi-omics profiling, including proteomics, lipidomics, and metabolomics, was carried out. Proteomics and metabolomics datasets were normalized using median normalization, whereas Lipidomics data were normalized using internal standards. All datasets were subsequently log₁₀-transformed. Differential analysis was independently conducted for each omics to assess the reproducibility and comparability of treatment-induced responses across platforms. For the volcano plots, statistical significance of individual features was determined using the t-test, followed by correction for multiple comparisons using the False Discovery Rate (FDR) approach^126^, to reduce the risk of false positives. Significantly altered proteins were further analyzed using enrichment analysis via STRING^121^ to explore associated biological pathways and interactions. For lipidomics, hierarchical clustering heatmaps were generated based on statistically significant lipids to visualize overarching trends within lipid subclasses and to facilitate biological interpretation. Before multivariate analysis, normalized datasets were autoscaled. Exploratory analysis was conducted to identify global trends and the underlying structure in the data. For Proteomics data, Principal Component Analysis (PCA) was applied to visualize clustering trends associated with treatment and culture conditions. For metabolomics and lipidomics datasets, each including both positive and negative ionization mode variables, a Common Dimensions (ComDim)^127^ approach was employed. This method enabled the joint analysis of multiple data blocks, facilitating the extraction of shared structures across ionization modes. For all the exploratory analysis, the score plots reported include 98% confidence ellipses, representing the multivariate region expected to contain 98% of the samples under the assumption of multivariate normality. Samples outside the confidence ellipse are considered potential outliers. The loadings plots (Supplementary Fig. 16) were used to identify the features contributing most to the observed group separation in the principal component space. Only the most informative variables, selected based on their variance contribution, were displayed and color-coded according to their p-values. For each feature, the p-value was calculated using the most appropriate univariate statistical test, selected based on the number of groups and the distribution of the data. Normality within each group was assessed using the Lilliefors test, and depending on the outcome, either parametric tests (t-test or ANOVA) or non-parametric alternatives (Mann–Whitney U test or Kruskal–Wallis test) were applied. All p-values were corrected for multiple testing using the FDR method.

### Functional enrichment analysis and network visualization

Genes (adjusted p-value < 0.01) and proteins (adjusted p-value < 0.05) differentially modulated after FOLFIRI treatment were subjected to analysis using Cytoscape software (ver. 3.10.3, Java ver. 17.0.5)^128^. The analysis was built to reveal functional enrichment of GO-Biological Process, KEGG and REACTOME-Reactions terms with ClueGO plug-in (ver. 2.5.10)^129^. Significant nodes were filtered for Bonferroni step down-corrected for significance (p-values < 0.01). Then Compound Spring Embedder (CoSE) layout was applied for visualization of the network. Identified terms and pathways were grouped into functionally related clusters using the Markov Cluster Algorithm (MCL) via Autoannotate (ver. 1.5.2)^130^ and WordCloud (ver. 3.1.4)^131^ plug-ins.

### Statistical analysis

Statistical analyses were performed using GraphPad Prism software (version v10.5.0). No particular statistical method was used to define the sample size and no specific hypothesis was tested. Sample size (n) values are provided in the relevant figures and supplementary figures as well as the specific tests used to determine statistical significance among groups. All data were analysed from at least 3 independent experiments unless stated otherwise. In the drug screening experiments, when necessary, the upper and lower boundaries for outliers removal was set at two standard deviations and never exceeded the 30% of the technical repetitions.

## Supplemental informations

**Supplementary Figure 1.**
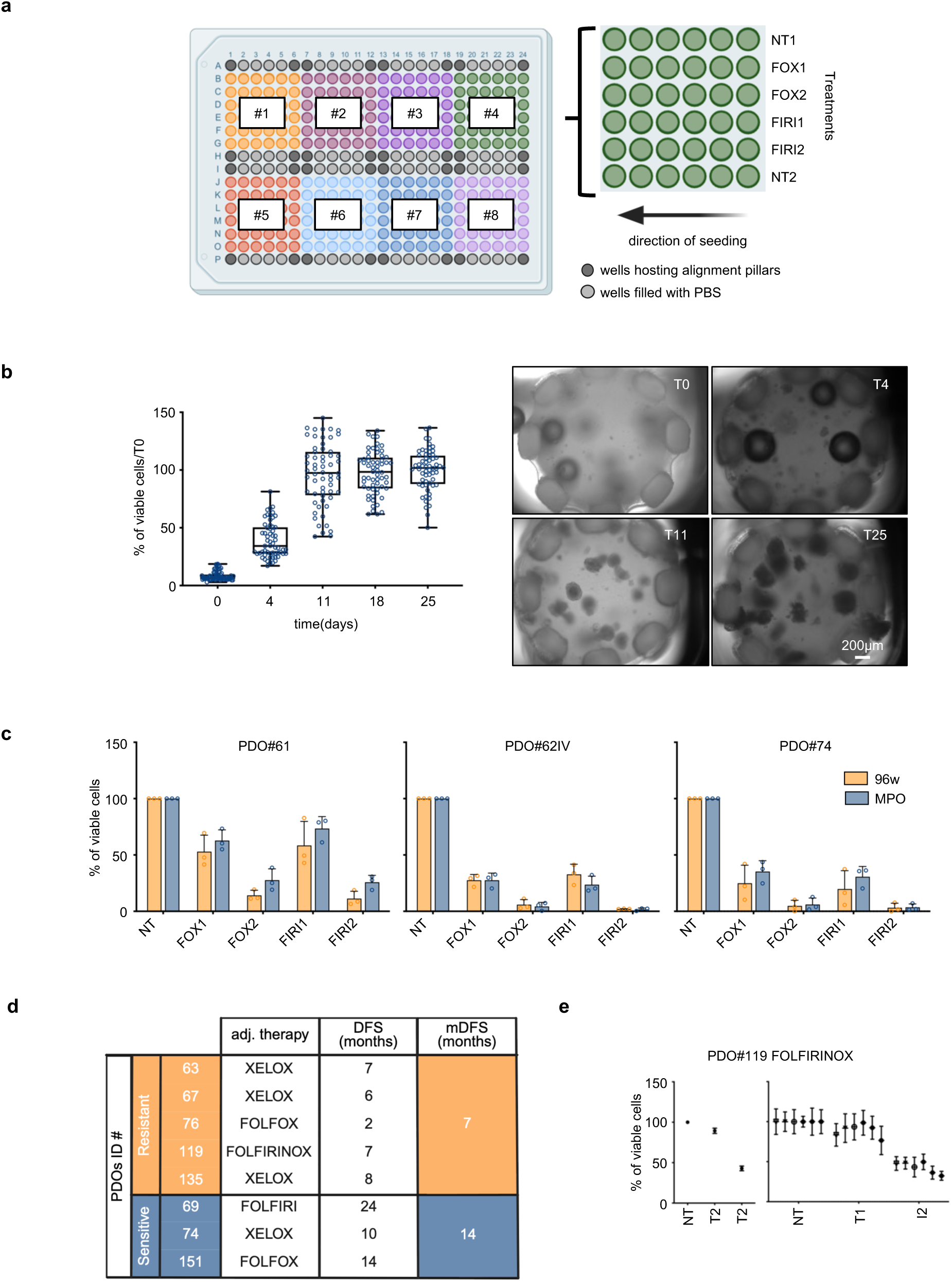
Drug sensitivity test on organoids grown and treated in the MPO. **a**. Schematic representation of a 384-well plate filled with 8 grids for a total of 288 single NESTs dipped in the culture medium. Different colours highlight the area encompassed by each grid. The empty wells hosting the alignment pillars or filled with PBS are depicted in dark and light grey, respectively. The direction of the seeding is also shown. A scheme of the treatment layout used in the drug sensitivity tests with six technical replicates for each treatment is provided. Organoids were exposed to FOLFOX (FOX) and FOLFIRI (FIRI), both used at two concentrations, one highest (FOX2 and FIRI2) and one 10–fold lower (FOX1 and FIRI1). Two vehicle-treated control lines were used in each grid, NT1 and NT2. One PDOs line was seeded in each grid. **b**. PDO #17 was cultured in the MPO and viability evaluated at different timepoints by CellTiter-Glo 3D assay and brightfield imaging analysis. Scale bar: 200 μm. Cell viability is reported as the percentage of viable cells at each time point relative to that at the time of seeding (T0). The culture was stopped at day 25 when no further increase in the growth was observed. Each dot refers to the measurement performed in one single pillar. For each dataset, boxes show top and bottom quartiles, centre line denotes the median value, and whiskers indicate minimum to maximum. Results of 7 independent experiments are reported (n= 59-77). **c**. A comparison between drug sensitivity tests on three PDOs lines performed in a conventional 96-well microplate setting or into the MPO is reported. For each condition, results are calculated as the mean percentage of viable cells in treated relative to untreated control samples (NT) averaged from three (in the 96-well microplate) or six technical repetitions. In each panel, the mean (± s.e.m.) of three independent experiments is reported for each PDOs line. A two-way ANOVA followed by Sidak’s multiple comparisons test was used to determine statistical significance between groups. **d**. In the table, the clinical outcome, measured as disease-free survival (DFS), of patients receiving adjuvant chemotherapy, was matched with the sensitivity response data of the corresponding organoids. Patients were treated with combinations of 5-FU and oxaliplatin or 5-FU and irinotecan regimens. The threshold to define sensitive and resistant PDOs was set as described in Fig.1c. The median DFS (mDFS) of both groups is shown. **e**. One patient received FOLFIRINOX adjuvant chemotherapy. The corresponding organoid line, PDO#119, was exposed to a combination of 5-FU, oxaliplatin and irinotecan (see Materials and Methods). Two concentrations were used, one highest (T2) and one 10–fold lower (T1). Plot on the left shows the sensitivities to the treatments reported as mean percentages of viable cells (mean ± s.e.m.) in treated relative to untreated control samples averaged from independent experiments (n= 6). The plot on the right shows the results of the independent experiments. For each condition results are calculated as the mean percentage (± s.d.) of viable cells in treated relative to untreated control samples (NT) averaged from technical repetitions.

**Supplementary Figure 2.**
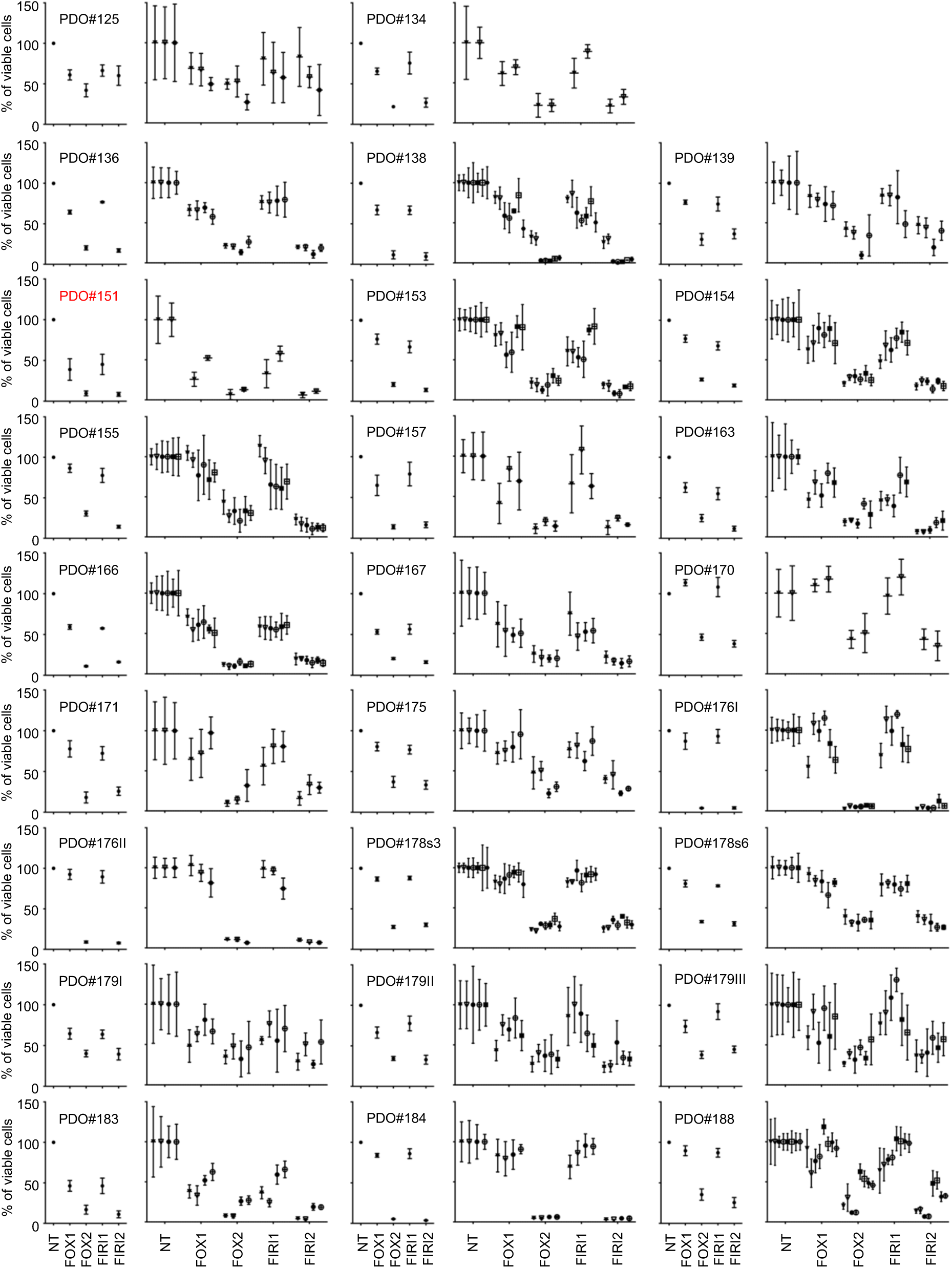
Prospective drug sensitivity test of organoids grown and treated in the MPO. A panel of 26 PDOs lines were seeded in the platform and exposed to a combination of drugs that replicated two chemotherapeutic regimens commonly used to treat patients with mCRC, FOLFOX (FOX) and FOLFIRI (FIRI). Two concentrations were used, one highest (FOX2 and FIRI2) and one 10–fold lower (FOX1 and FIRI1). For each PDOs line, plots on the left show the sensitivities to the treatments reported as mean percentages of viable cells (mean ± s.e.m.) in treated relative to untreated control samples (NT) averaged from independent experiments. Plots on the right show the results of the independent experiments. For each condition results are calculated as the mean percentage (± s.d.) of viable cells in treated relative to untreated control samples (NT) averaged from technical repetitions. In red organoids derived from patients receiving adjuvant chemotherapy.

**Supplementary Figure 3.**
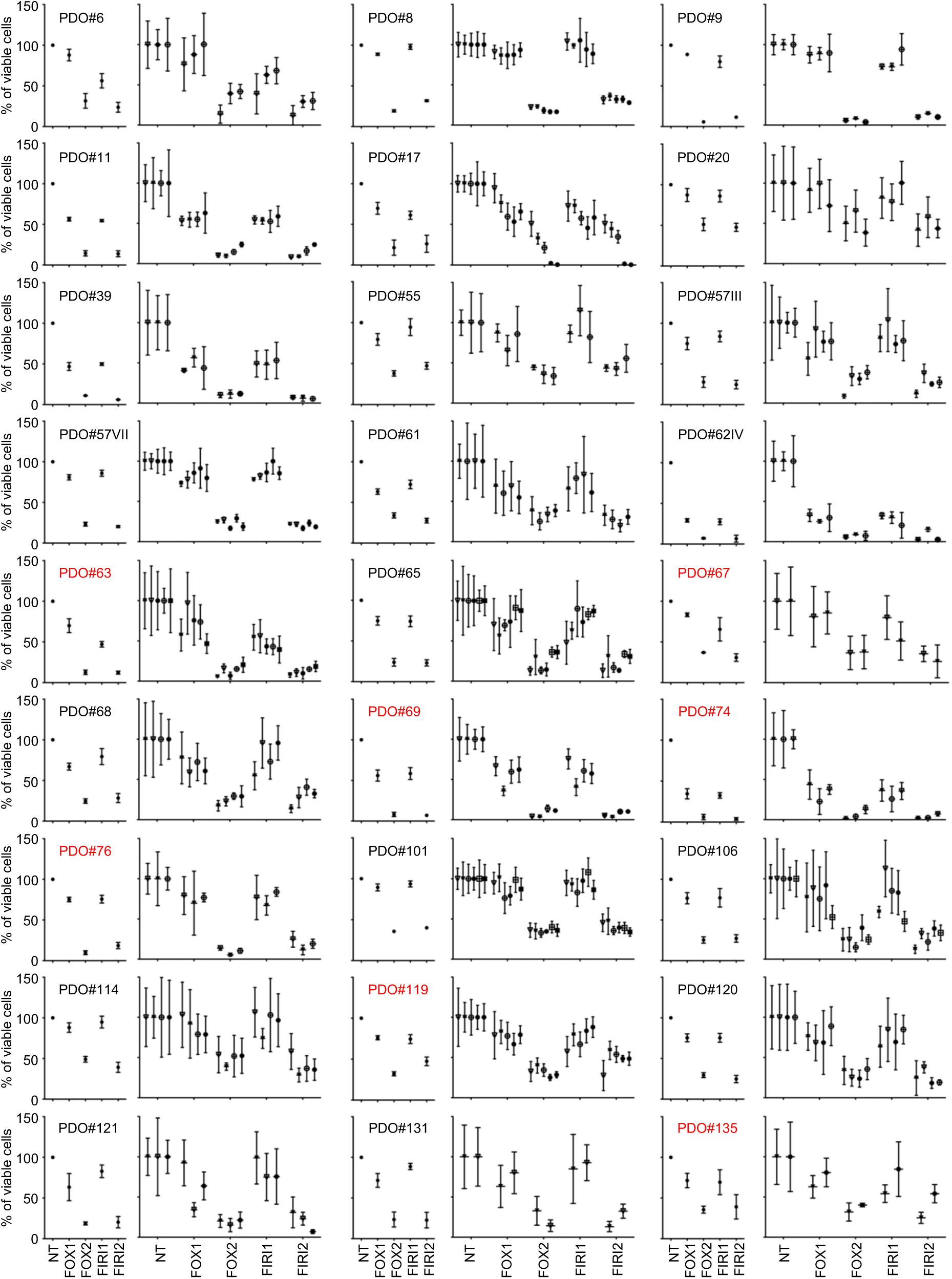
Retrospective drug sensitivity test of organoids grown and treated in the MPO. A panel of 27 PDOs lines were seeded in the platform and exposed to a combination of drugs that replicated two chemotherapeutic regimens commonly used to treat patients with mCRC, FOLFOX (FOX) and FOLFIRI (FIRI). Two concentrations were used, one highest (FOX2 and FIRI2) and one 10–fold lower (FOX1 and FIRI1). For each PDOs line, plots on the left show the sensitivities to the treatments reported as mean percentages of viable cells (mean ± s.e.m.) in treated relative to untreated control samples averaged from independent experiments. Plots on the right show the results of the independent experiments. For each condition results are calculated as the mean percentage (± s.d.) of viable cells in treated relative to untreated control samples (NT) averaged from technical repetitions. Based on the sensitivity to the lower doses, the threshold that distinguished the two groups resulted in a 40% reduction of viability relative to the untreated control. Most of the PDOs were either sensitive or resistant to both treatments although there were two exceptions: PDO#63 which was more sensitive to FOLFIRI and PDO#163, which was more sensitive to FOLFOX. In red organoids derived from patients receiving adjuvant chemotherapy after surgery.

**Supplementary Figure 4.**
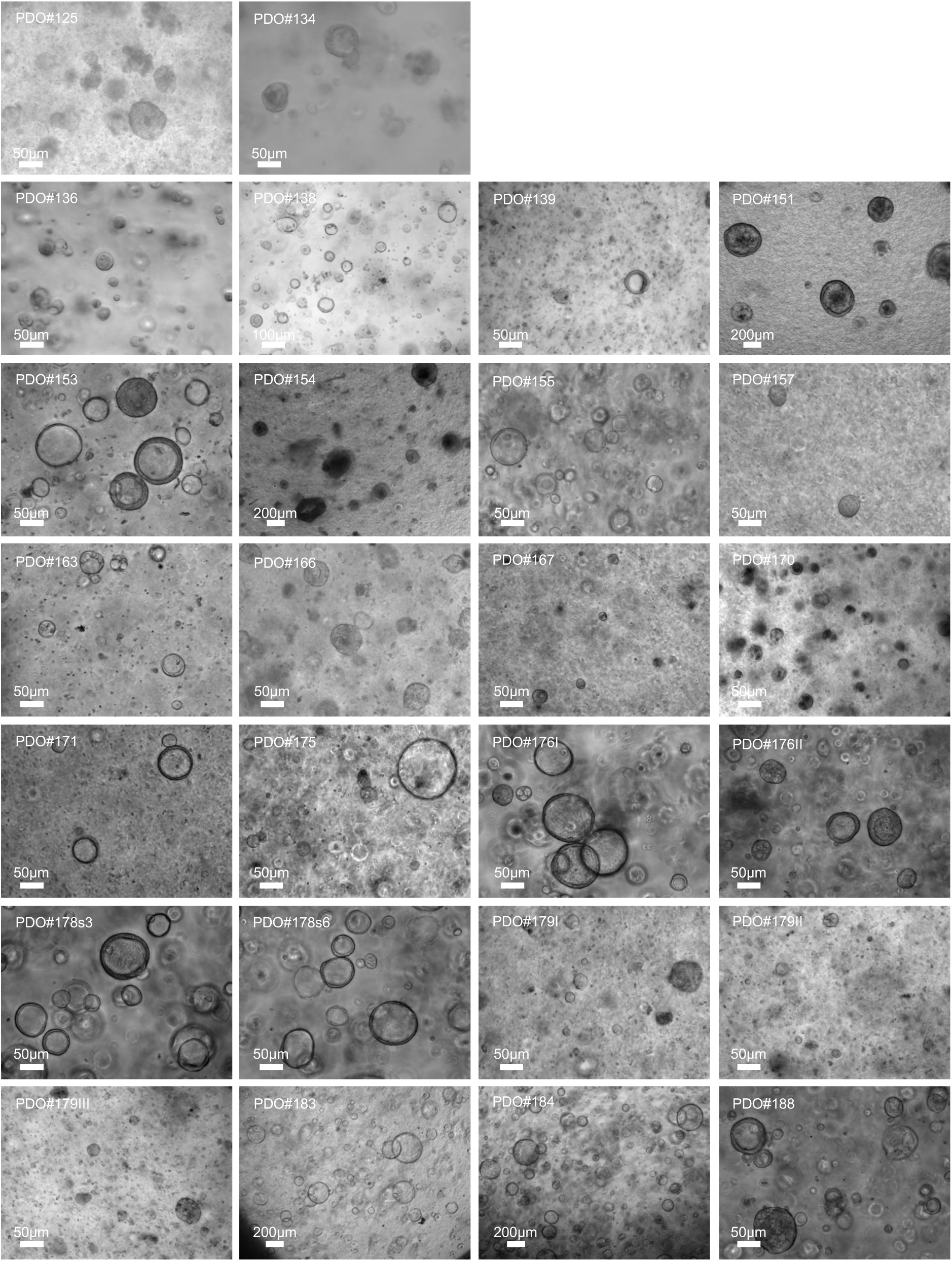
Representative brightfield images of the PDOs lines used for the prospective study. Images were taken at the time of the first drug test in the MPO. The scale bar is shown in each image.

**Supplementary Figure 5.**
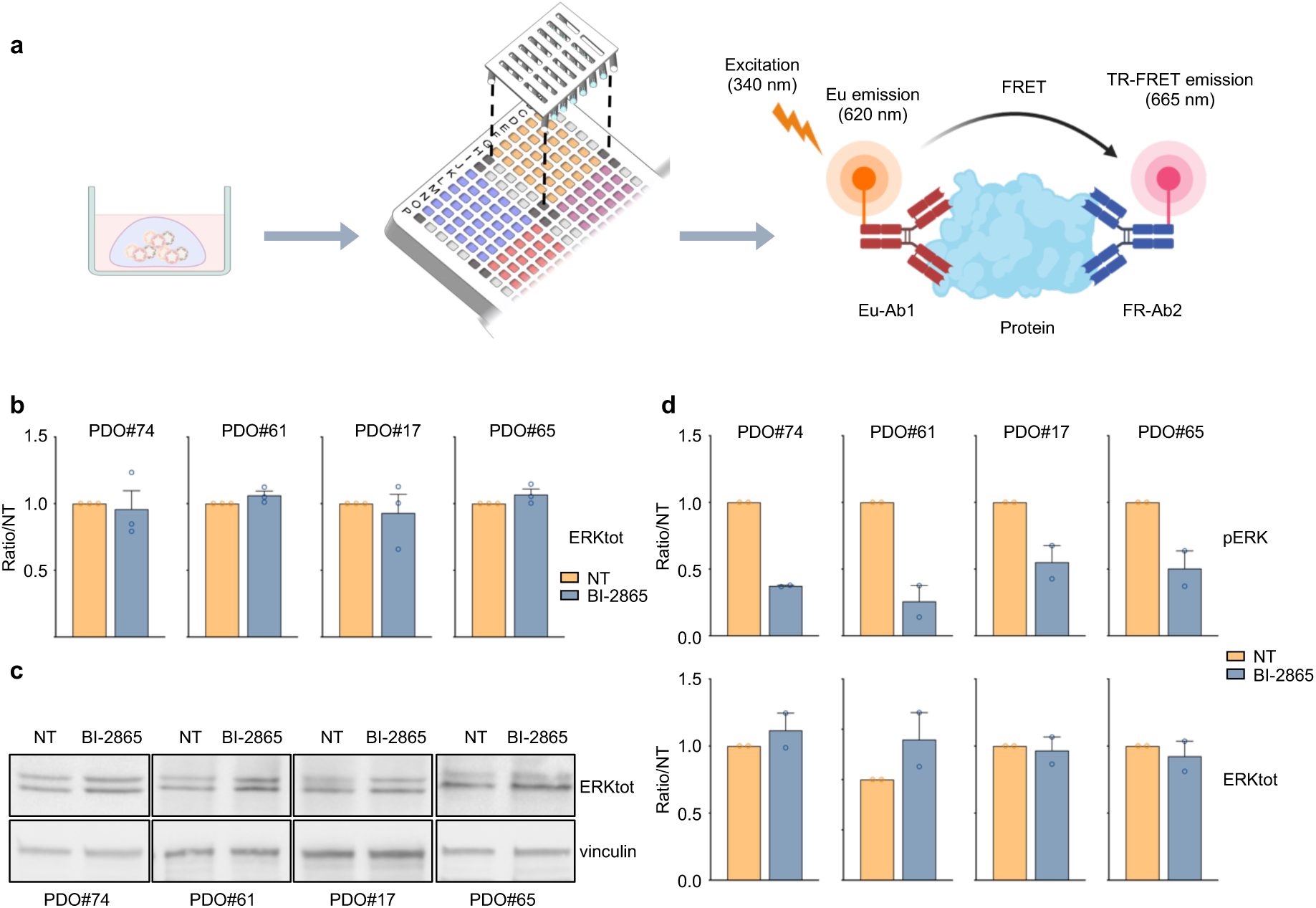
Target engagement by HTRF-based assays on organoids grown and treated in the MPO. **a**. Schematic representation of the experimental procedure for the HTRF-based assays in the MPO. **b**. A panel of four KRASmut PDOs lines were seeded in the platform and treated for 4 hours either with DMSO (NT) or BI-2865 (KRASi). Levels of total ERK (ERKtot) were assessed in HRTF-based assays. Results are reported as the levels of the protein in the treated samples relative to the untreated counterpart (ratio/NT) averaged (± s.e.m.) from three independent experiments, each performed in at least three technical replicates per condition. **c**. Western blot analysis for the expression of ERKtot was performed on the same PDOs lines grown and treated as in (b) but not in a miniaturized setting. Vinculin was used as loading control. One representative experiment of two is shown. **d**. Quantitative analysis of the western blot presented in (c) and in Fig. 2b. Results are reported as the levels of each protein (normalized to the loading control) in the treated samples relative to the untreated counterpart (ratio/NT) averaged (± s.e.m.) from two independent experiments.

**Supplementary Figure 6.**
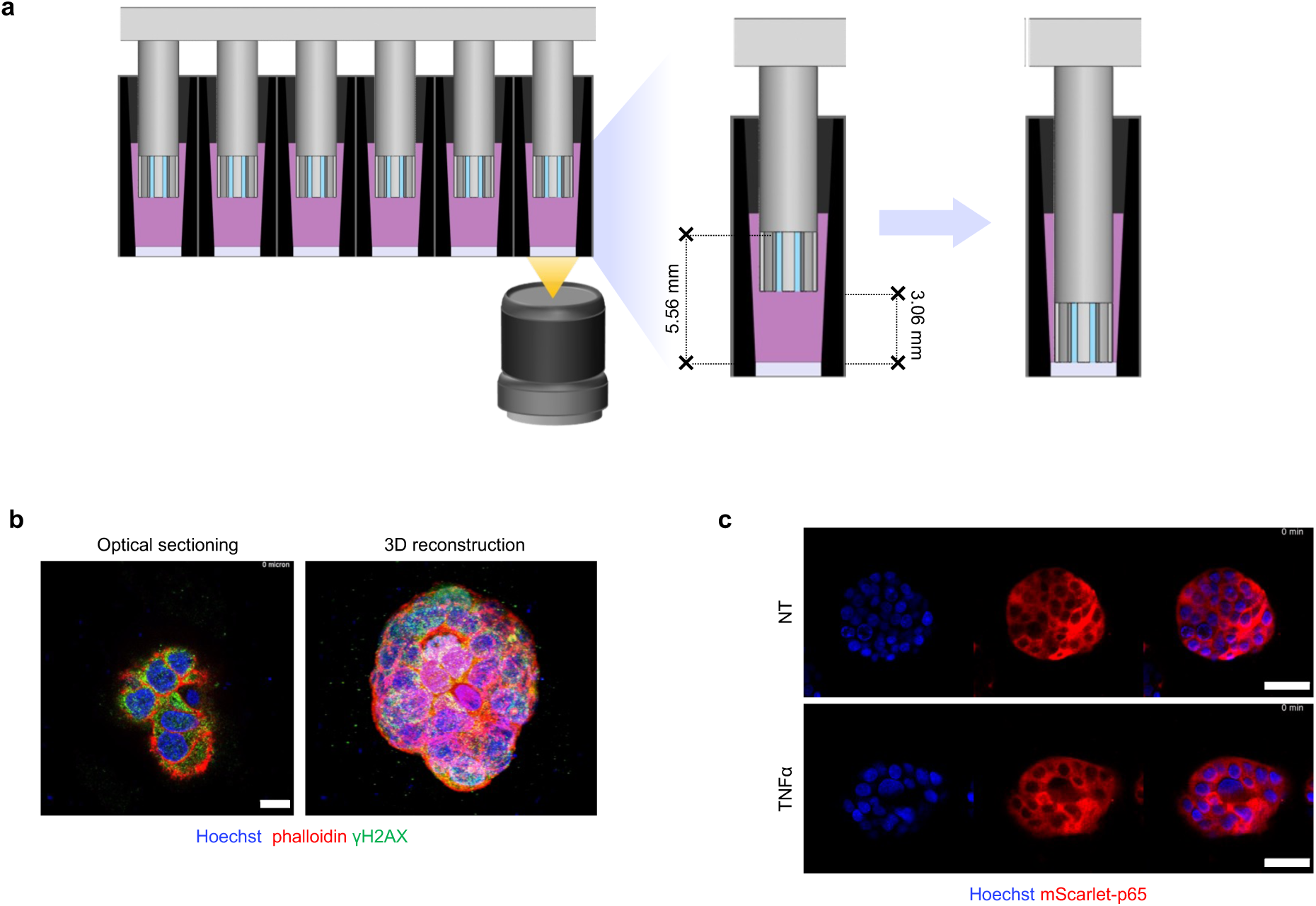
High-content imaging of PDOs cultured and stained in the MPO. **a**. For confocal imaging purposes, the NESTs grid architecture was modified to position the surface of the matrix embedding the PDOs closer to the bottom of the wells. **b**. Snapshots of representative movie of optical sectioning (left, related to Supplementary movie 1) and 3D reconstruction (right, related to Supplementary movie 2) of immunofluorescence of PDO#6 grown in the MPO, fixed and stained for Hoechst, phalloidin and γH2AX. Scale bar: 10 μm **c**. Snapshots of time-lapse movies (related to Supplementary movie 3 and 4) showing an optical section of spheroids derived from MCF7 cells expressing mScarlet-p65 and stained with Hoechst, and then left untreated (NT) or treated with TNFα (TNFα). The individual channels and merged channels are shown. Scale bar: 50 μm.

**Supplementary Figure 7.**
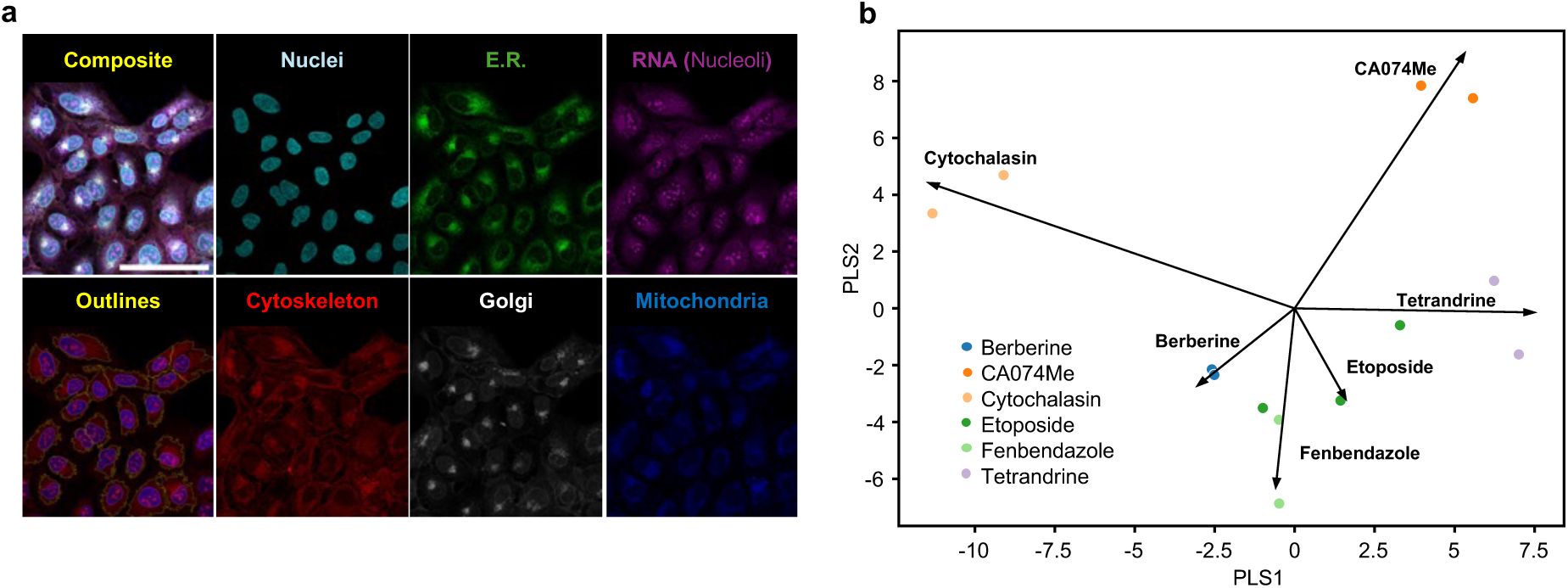
Cell Painting in U2OS cells. U2OS cells were left untreated or treated for 48 hours with Tetrandrine, Fenbendazole, Ca074Me, Etoposide, Cytochalasin B and Berberine (see Material and Methods). **a**. Individual fluorescent channels of a representative single-plane confocal image. A composite image of all channels and segmentation-derived outlines are shown. Scale bar: 100 μm. **b**. The biplot shows the results of PLS analysis. The profiles of multiple Cell Painting experiments in the PLS space were fitted to maximize the difference between treatments. The arrows represent the loading vectors for each drug and indicate the relative contribution to each PLS coordinate displayed.

**Supplementary Figure 8.**
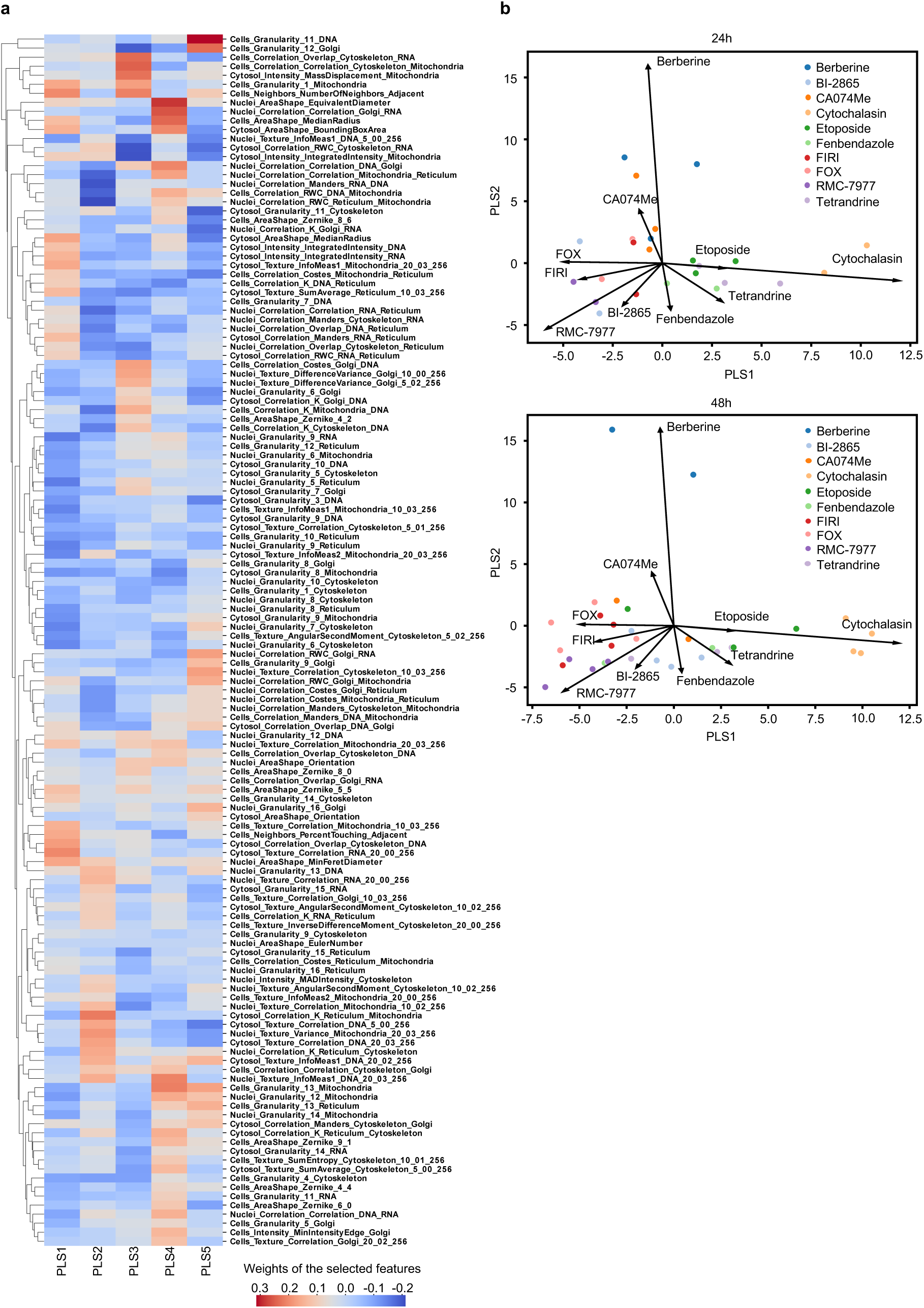
Cell Painting on organoids grown and treated in the MPO. PDO#17 was seeded in the MPO and treated as reported in Fig. 3. **a**. The heatmap shows the coefficient of each Cell Painting feature on each component in PLS regression. Red and blue colors indicate, respectively, the positive or the negative association of a feature with the PLS component. Hierarchical clustering of features was performed on the distance matrix resulting from the computation of pairwise correlations between features vectors. **b**. The biplots show the results of PLS analysis. Each plot is a subset of samples at different times, as displayed in Fig. 3. The plot represents the profiles of multiple Cell Painting experiments in the PLS space fitted to maximize the difference between treatments. The arrows represent the loading vectors for each drug and indicate the relative contribution to each PLS coordinate displayed.

**Supplementary Figure 9.**
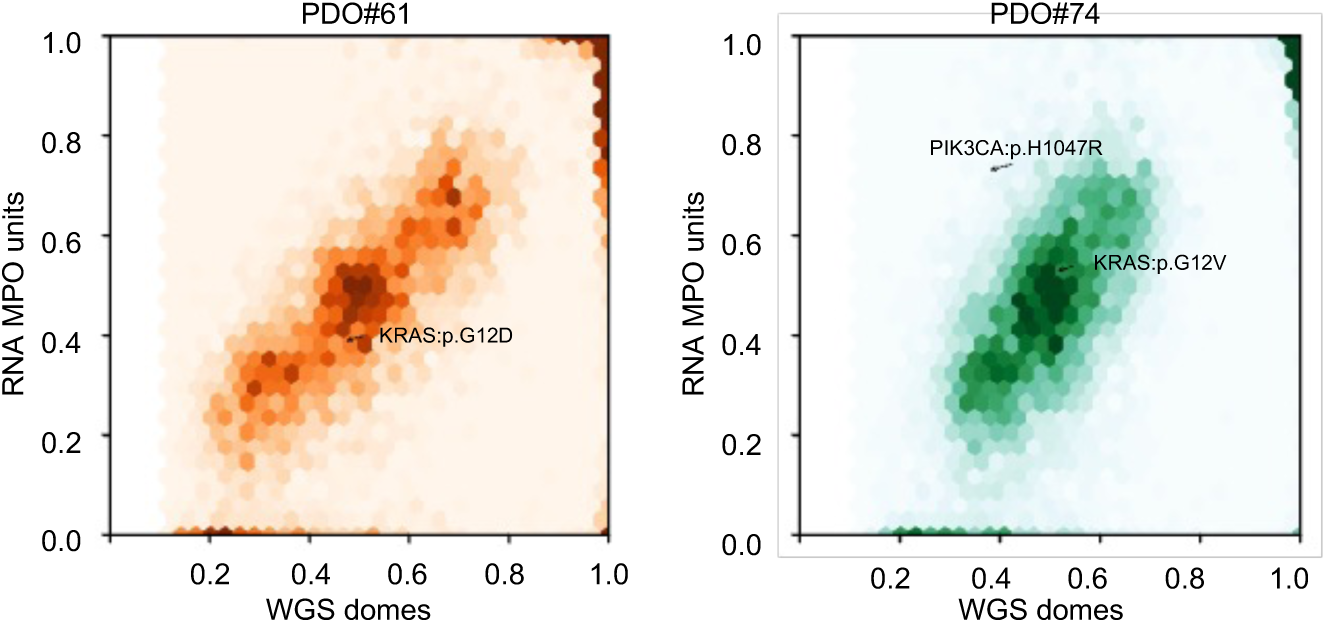
MPO enables the integration of multiple genomics platforms. Hexbin-plots showing comparative Variant Allele Frequencies (VAFs) between high coverage WGS and RNA-seq on pillars showing somatic mutations for PDO#61 and PDO#74. Driver mutations are highlighted.

**Supplementary Figure 10.**
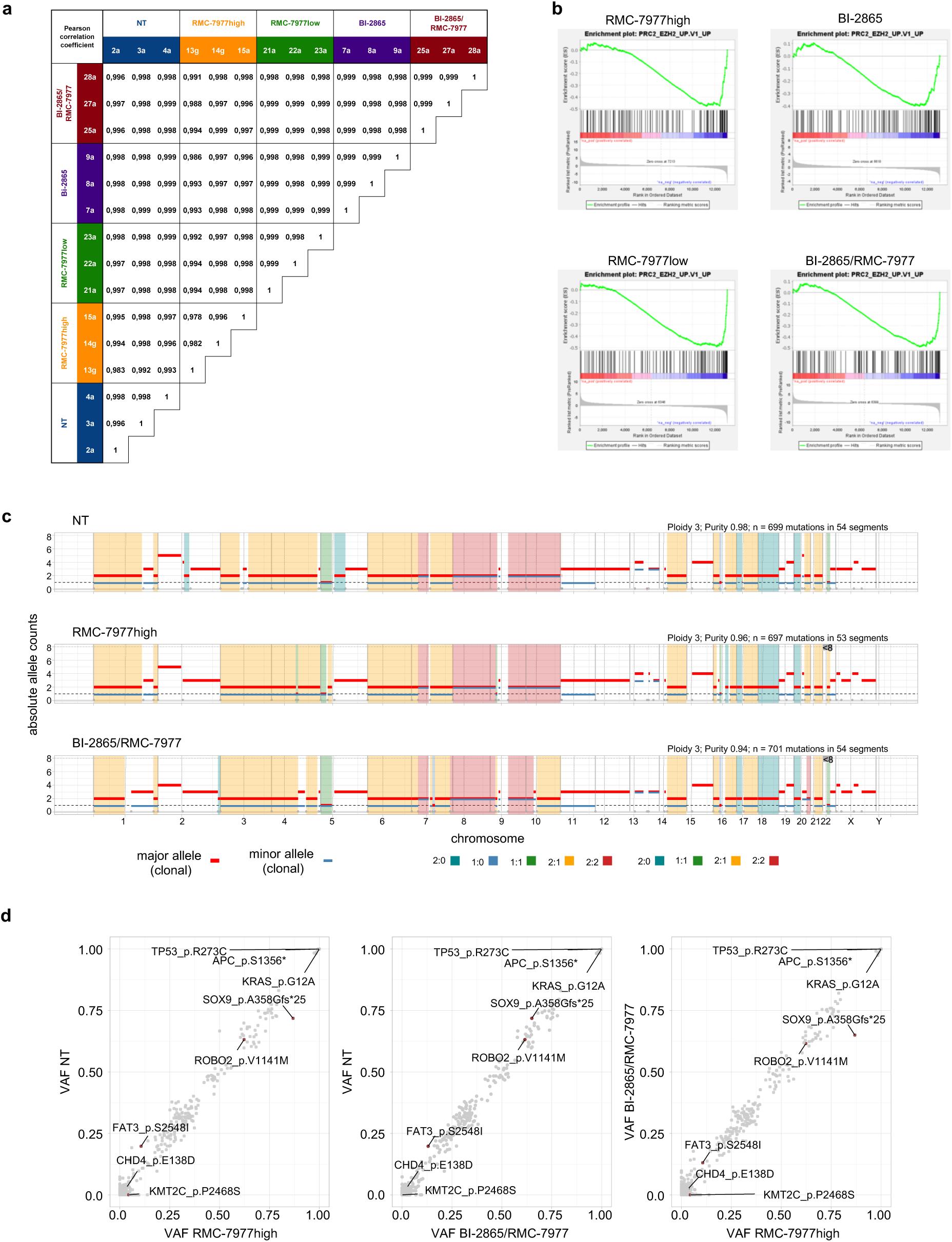
Modelling development of drug resistance in the MPO. PDO#17 was seeded in the MPO, treated and analysed as reported in Fig. 5a and b. **a**. The table displays the Pearson correlation coefficient for the comparisons between low-pass WGS. **b**. The GSEA leading edge plots, as obtained after sequence analysis of RNA, depict the repression of PRC2–EZH2 regulated signature in treated vs untreated organoids. Of note, BI-2865-treated organoids presented the same enrichment profile of the other treated organoids even if it was below the threshold of significance (FDR q-val= 0.0134). **c**. Genome wide copy number profile showing the absolute allele counts across all chromosomes for the reported treatments was performed with ASCAT. The analysis also provided estimates of tumor ploidy and purity. **d**. Scatter-plots comparing the variant allele frequency (VAF) of somatic mutations in treated vs untreated samples (NT) and between the two treatment groups. Each dot represents a single mutation (grey dots no drivers, red dots drivers).

**Supplementary Figure 11.**
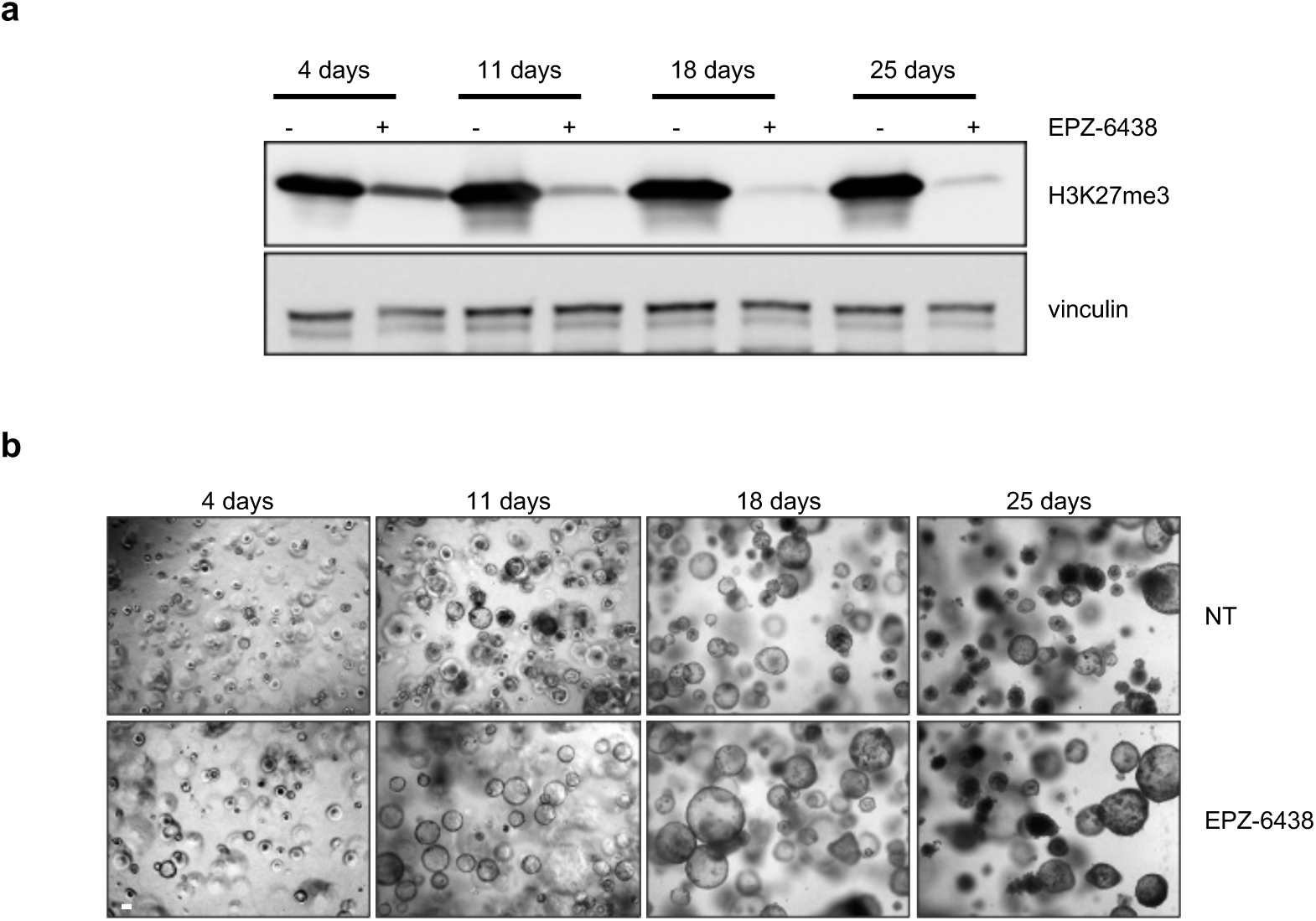
EZH2 target inhibition. PDO #17 was left untreated (NT) or treated with EP-6438. **a**. At the reported time points, organoids were collected for western blot analysis of H3K27me3 levels. Vinculin was used as loading control. One representative experiment of two is shown. **b**. Representative brightfield images of NT or EP-6438-treated organoids. Scale bar: 50 μm.

**Supplementary Figure 12.**
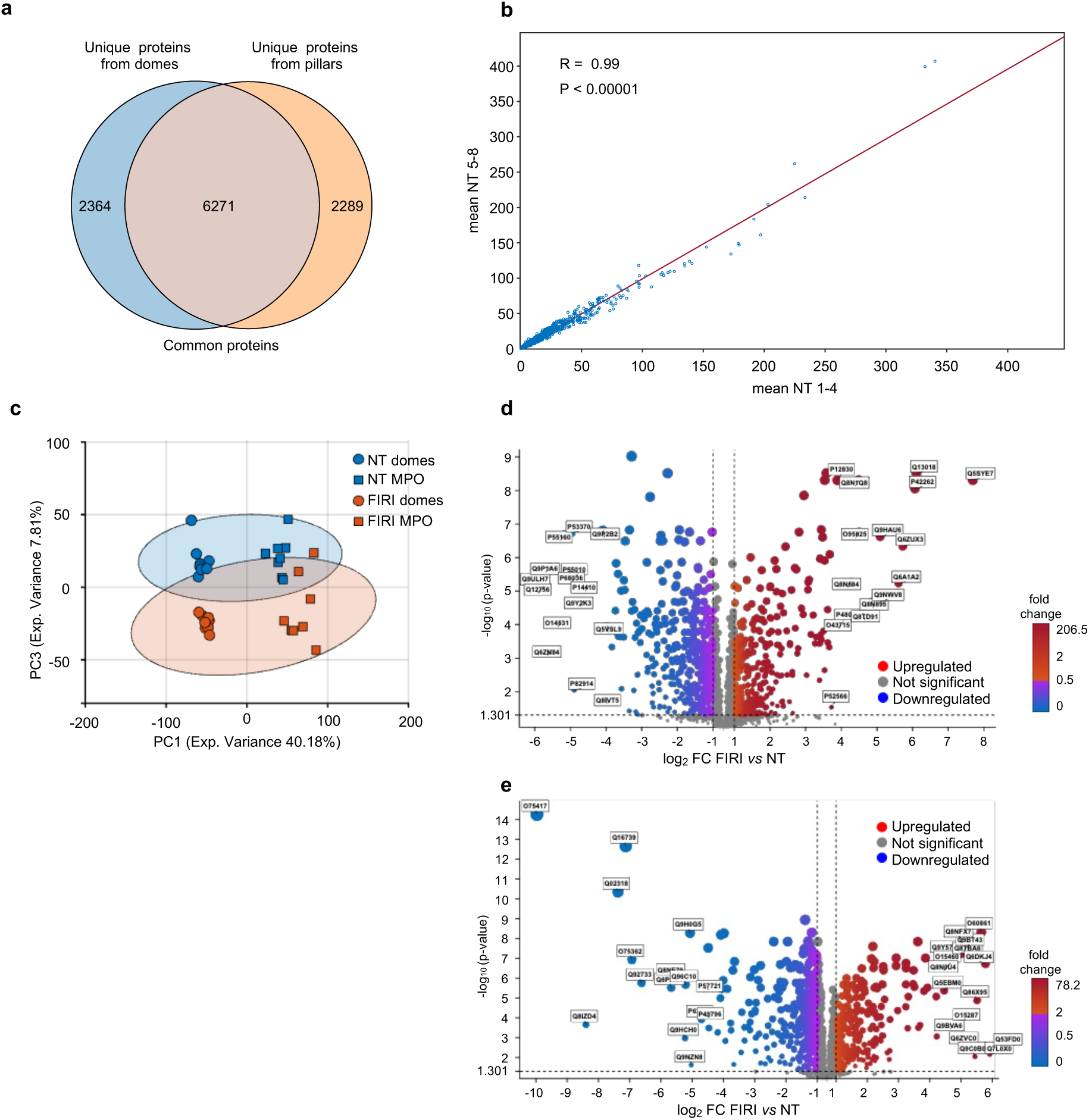
Proteomics analysis on organoids grown and treated in the MPO. PDO#61 was seeded in the MPO and left untreated (NT) or treated with FOLFIRI (FIRI) either in the MPO or in conventional conditions (domes) (n=8 for each experimental setting). At the end of the treatment, organoids were recovered and prepared for proteomic analysis. **a**. The Venn diagrams report the number of unique or shared proteins identified in the MPO or in the conventional culture (domes). **b**. Pearson’ correlation plot calculated between the mean of 1-4 and 5-8 pillars of MPO-grown NT samples. **c**. Score plot (PC1 vs PC3) from PCA of the proteomic datasets reporting the clustering of NT and FOLFIRI-treated organoids in the two experimental settings. **d-e.** Volcano plots depict differentially FOLFIRI-regulated proteins obtained from organoids grown in the MPO (d) or in conventional conditions (e). Statistical significance was determined using the t-test followed by Benjamini-Hochberg FDR procedure. The most significant proteins are annotated. Down-regulated (log2(fold change)<1 and p-value<0.05) proteins in FOLFIRI-treated samples are shown in blue, while up-regulated (log2(fold change)>1 and p-value<0.05) proteins are shown in red.

**Supplementary Figure 13.**
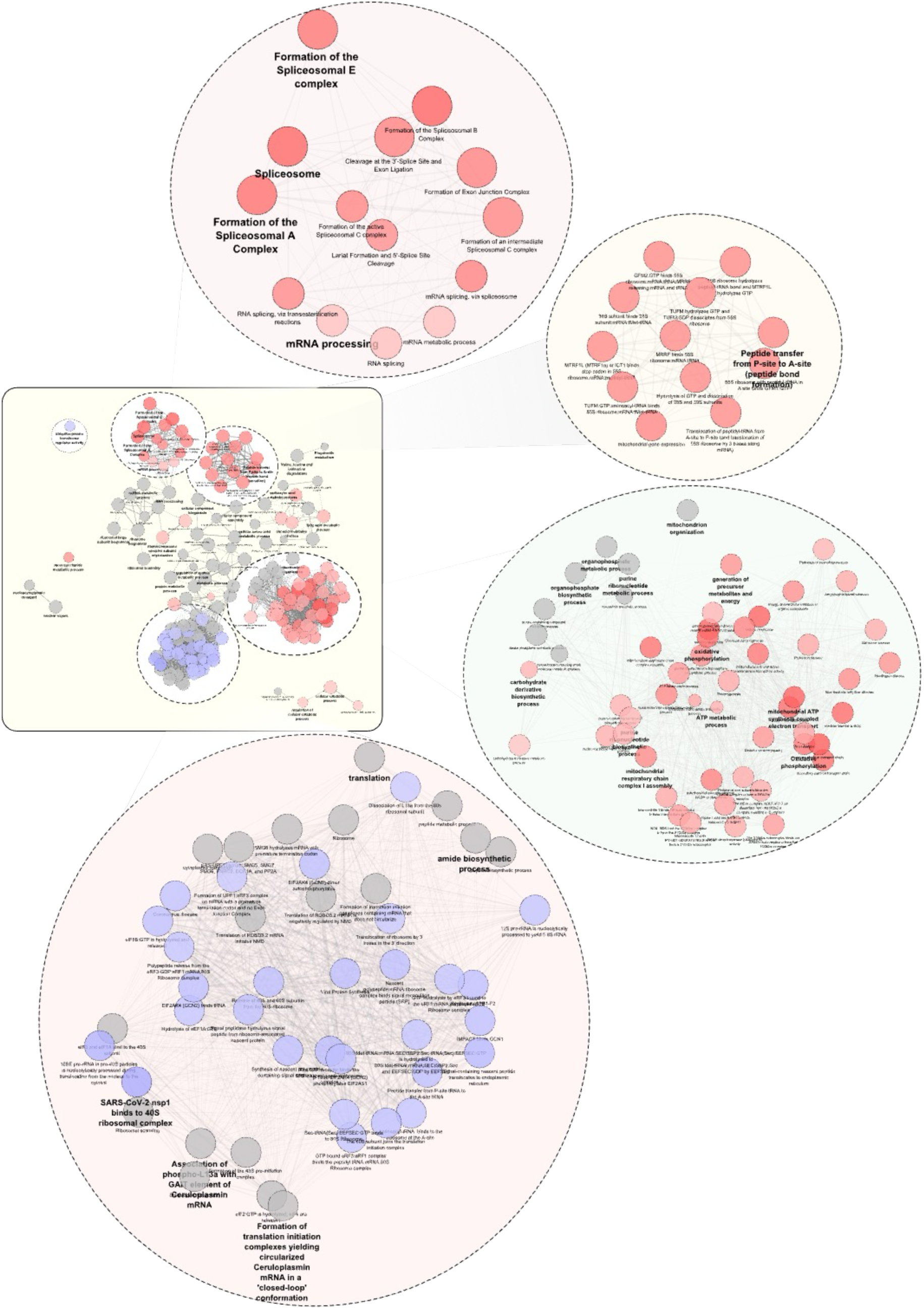
(related to. Figure 6c). Complete visualization of the network generated on the functional enrichment analysis of RNA and proteins quantified as described in Materials and Methods. Enriched terms are displayed as single nodes, edges denote overlapping of genes in the linked nodes. Balloons indicate zoom-ins in selected clusters. Gene sets with a FDR q-value < 0.01 were considered significant.

**Supplementary Figure 14.**
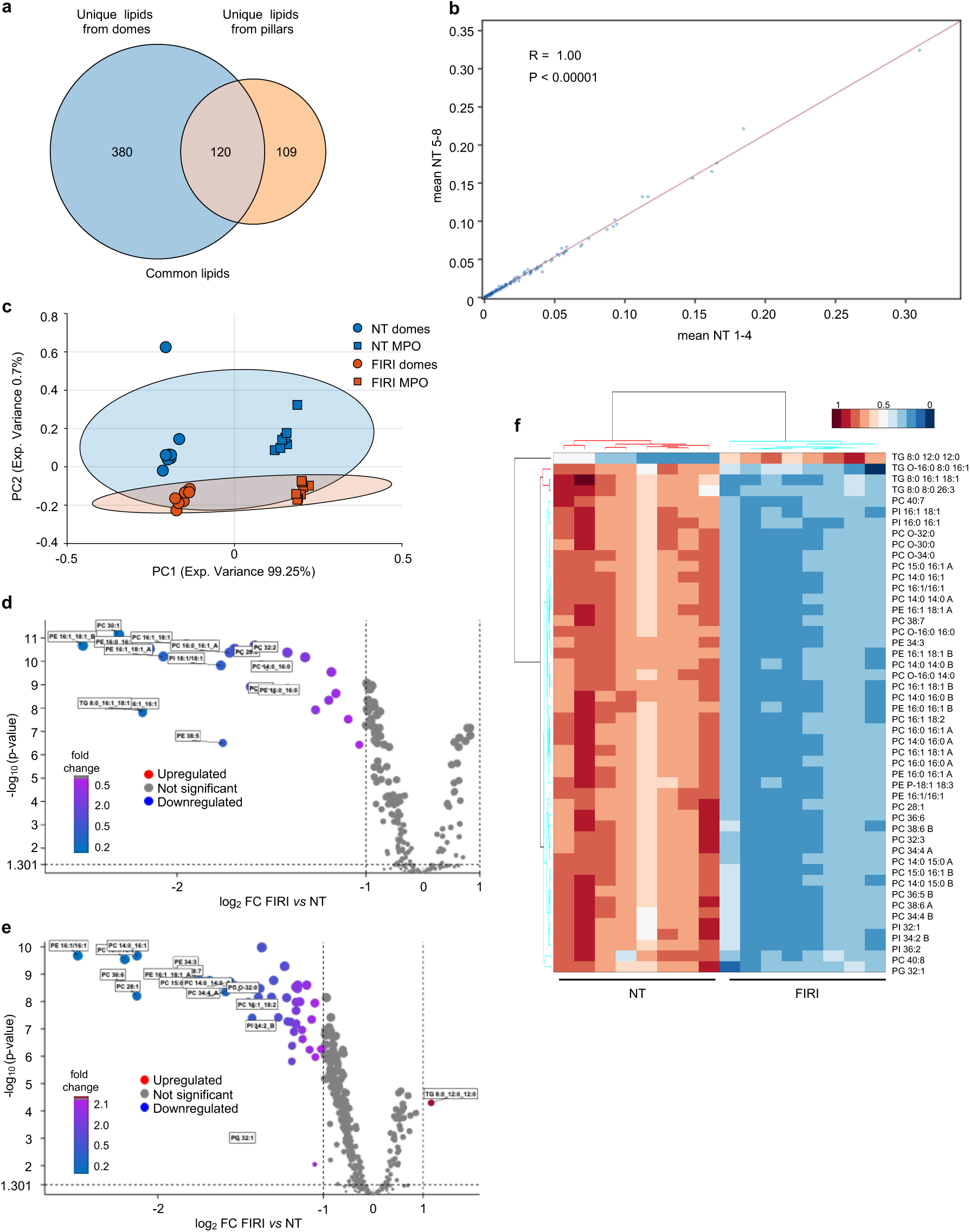
Lipidomics analysis on organoids grown and treated in the MPO. PDO#61 was seeded in the MPO and left untreated (NT) or treated with FOLFIRI (FIRI) either in the MPO or in conventional conditions (domes) (n=8 for each experimental setting). At the end of the treatment, organoids were recovered and prepared for lipidomics analysis. **a**. The Venn diagrams report the number of unique or shared lipids identified in the MPO or in the conventional culture (domes). **b**. Pearson’ correlation plot calculated between the mean of 1-4 and 5-8 pillars of MPO-grown NT samples. **c**. Score plot (PC1 *vs* PC2) from PCA of the lipidomics datasets reporting the clustering of NT and FOLFIRI-treated organoids in the two experimental settings. **d-e**. Volcano plots depict differentially FOLFIRI-regulated lipids obtained from organoids grown in the MPO (d) or in conventional conditions (e). Statistical significance was determined using the t-test followed by Benjamini-Hochberg FDR procedure. The most significant lipids are annotated. Down-regulated (log2(fold change)<1 and p-value<0.05) lipids in FOLFIRI-treated samples are shown in blue, while up-regulated (log2(fold change)>1 and p-value<0.05) lipids are shown in red. **f**. The heatmap depicts the results of lipidomics analysis performed in NT and FOLFIRI-treated organoids grown in the conventional culture (domes). The variables shown were selected based on significance criteria from the corresponding volcano plot.

**Supplementary Figure 15.**
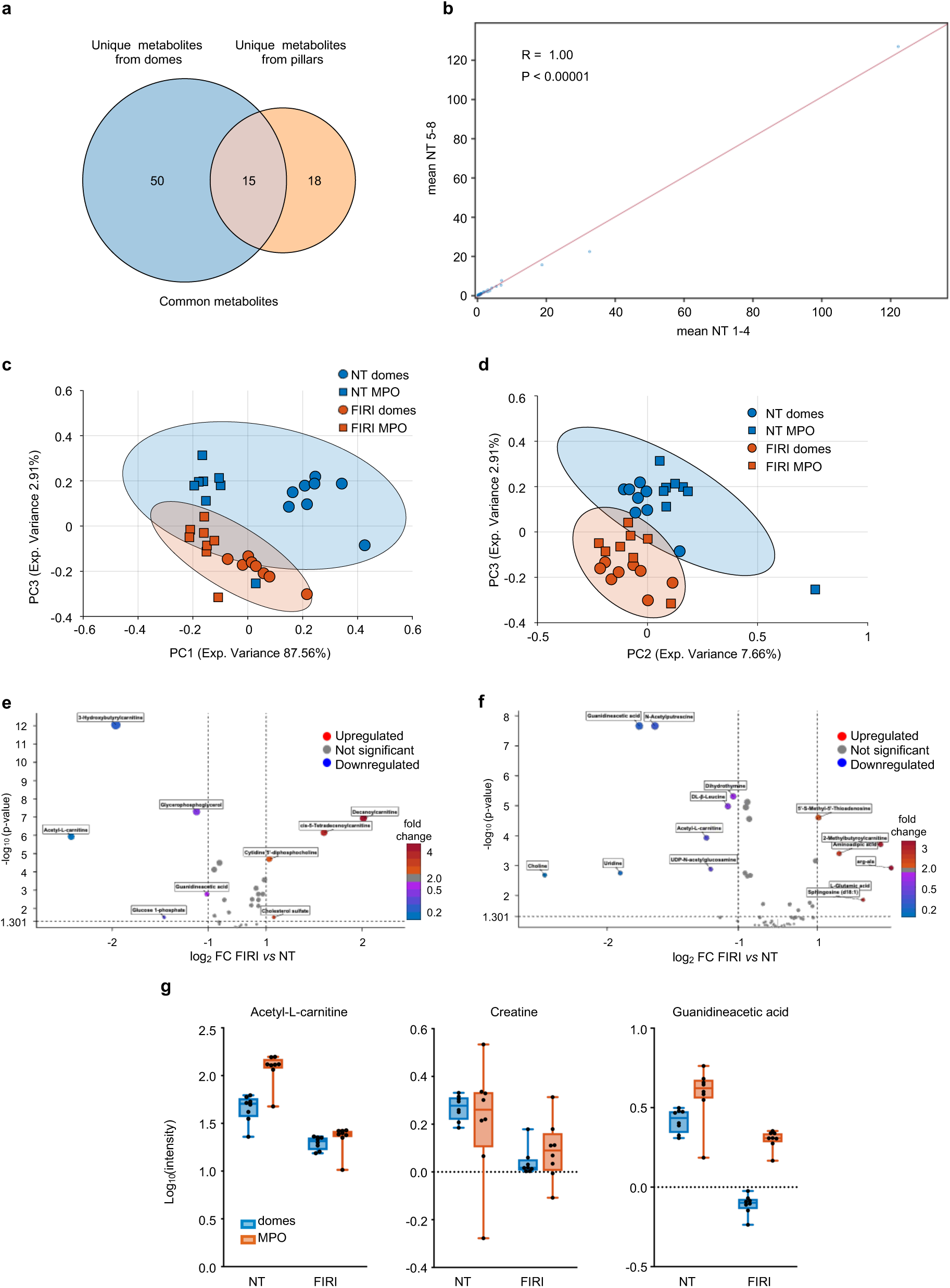
Metabolomics analysis on organoids grown and treated in the MPO. PDO#61 was seeded in the MPO and left untreated (NT) or treated with FOLFIRI (FIRI) either in the MPO or in conventional conditions (domes) (n=8 for each experimental setting). At the end of the treatment, organoids were recovered and prepared for metabolomic analysis. **a**. The Venn diagrams report the number of unique or shared metabolites identified in the MPO or in the conventional culture (domes). **b**. Pearson’ correlation plot calculated between the mean of 1-4 and 5-8 pillars of MPO-grown NT samples. **c-d**. Score plots from PCA of the metabolomics datasets reporting the clustering of NT and FOLFIRI-treated organoids in the two experimental settings along PC1 and PC3 (c) or PC2 and PC3 (d). **e-f**. Volcano plots depict differentially FOLFIRI-regulated metabolites obtained from organoids grown in the MPO (d) or in conventional conditions (e). Statistical significance was determined using the t-test followed by Benjamini-Hochberg FDR procedure. The most significant metabolites are annotated. Down-regulated (log2(fold change)<1 and p-value<0.05) metabolites in FOLFIRI-treated samples are shown in blue, while up-regulated (log2(fold change)>1 and p-value<0.05) metabolites are shown in red. **g**. Plots report the distribution of Acetyl-N-carnitine, Creatine and Guanidineacetic acid in NT and FOLFIRI-treated organoids in the two experimental settings. For each dataset, boxes show top and bottom quartiles, centre line denotes the median value, and whiskers indicate minimum to maximum.

**Supplementary Figure 16.**
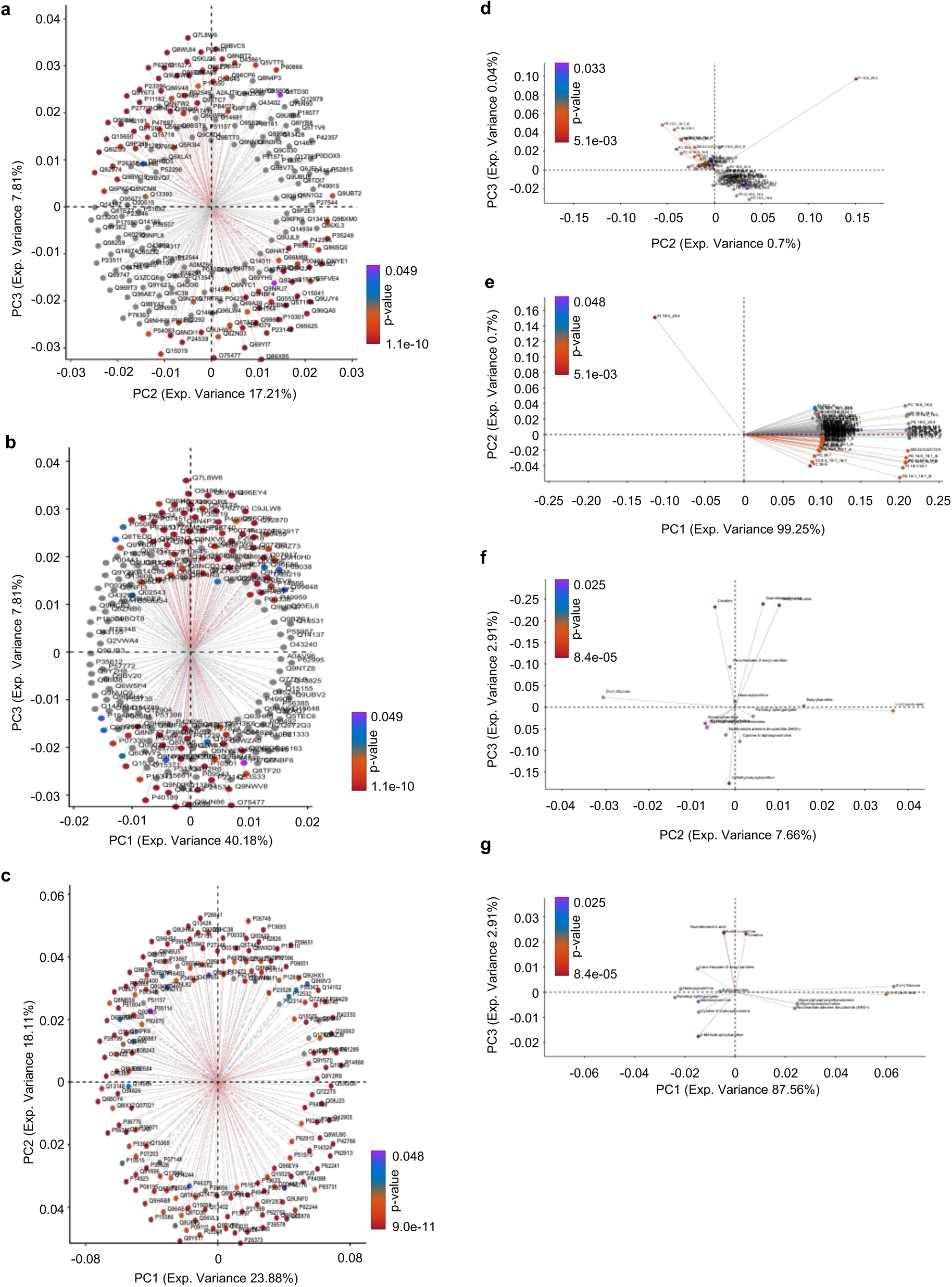
Loadings plots for the MS-multi omics exploratory analyses on organoids. Loadings plots highlight the variables most responsible for groups separation in the multivariate space along PCs, displaying those with higher contribution to variance, coloured according to their p-values. **a-c**. Loadings plots of most relevant proteins of the proteomics datasets relative to PCA in Fig. 6a (a), Supplementary Fig. 12c (b) and Fig. 6d (c). **d-e**. Loadings plots of most relevant lipids of the lipidomics datasets relative to COMDIM in Fig. 6d (d), Supplementary Fig. 14c (e). **f-g**. Loadings plots of most relevant metabolites of the metabolomics datasets relative to COMDIM in Supplementary Fig. 15c (f) and Supplementary Fig. 15d (g).

**Supplementary Table 1.**

Clustering and Mutual Information (MI) calculation for all features as obtained by analysis of features extracted by images of PDOs along with PLS loadings relative to the features indicated in each sheet. For both PDOs and U2OS cells, allfeatures and filteredfeatures’ loadings are reported respectively.

**Supplementary Table 2.**

Results and metadata for the functional enrichment analysis performed via ClueGO within Cytoscape. Modulated genes as seen by transcriptomic analysis and proteomic analysis are defined as Cluster #1 and Cluster #2, respectively. Resulting nodes and edges attributes are reported with the corresponding relative Kappa Scores.

**Supplementary Table 3.**

PDO#17 was seeded in the MPO, treated and analysed as reported in Fig. 5a and b. In the table the cancer-relevant pathways modulated upon drug exposure as obtained by GSEA after sequence analysis of RNA.

**Supplementary movie 1.** Optical sectioning relative to the snapshot image in Supplementary Fig. 6b.

**Supplementary movie 2.** 3D reconstruction relative to the snapshot image in Supplementary Fig. 6b.

**Supplementary movie 3.** Time-lapse movie relative to the snapshot image of untreated (NT) MCF7 spheroids in Fig. 2f and Supplementary Fig. 6c.

**Supplementary movie 4.** Time-lapse movie relative to the snapshot image of TNFα-treated MCF7 spheroids in Fig. 2f and Supplementary Fig. 6c.

